# Insights from pooled CRISPRi single-cell screens in K562 cells reveal gene functions, regulatory networks, and highlight opportunities and limitations

**DOI:** 10.1101/2025.10.23.684112

**Authors:** Hong Zhang, Peifen Zhang, Eric Bindels, Eskeatnaf Mulugeta

## Abstract

Pooled CRISPR screening combined with single-cell RNA sequencing (scRNA-seq) has emerged as a powerful strategy for dissecting gene function and reconstructing gene regulatory networks (GRNs) in complex biological systems. This approach enables high-throughput, parallel perturbation of multiple genes while providing transcriptome-wide readouts at single-cell resolution, overcoming many limitations of traditional arrayed screens. However, its broader application remains limited by technical challenges, including variable perturbation efficiency and difficulties in accurately identifying perturbed cells.

In this study, we adapted and applied a modified CRISPR droplet sequencing (CROP-seq) protocol using CRISPR interference (CRISPRi) in K562 cells to knockdown six transcription factors (TFs): LMO2, TCF3, LDB1, MYB, GATA2, and RUNX1. Our modified approach, which allows direct capture of sgRNAs from the cDNA library without a separate enrichment step, significantly improved sgRNA assignment per cell. We successfully achieved reproducible knockdown of three TFs (MYB, GATA2, and LMO2), captured the impact of these perturbations on the TF target genes, and enabled us to reconstruct their GRNs and identify key regulons and transcriptional targets. These networks revealed both previously established (such as LMO2 GATA2 interaction) and novel regulatory interactions, which we independently validated, providing new insights into hematopoietic transcriptional control. To assess the efficiency of CRISPRi based pooled perturbation, we additionally analyzed publicly available pertrub-seq CRISPRi datasets and found that only ∼40–50% of targeted genes led to effective knockdown, underscoring the variability in perturbation efficiency across experiments.

Together, our results demonstrate both the potential and the current technical limitations of pooled CRISPRi-based single-cell screens. While this integrated approach holds great promise for high-resolution functional genomics, further optimization and standardized benchmarking are essential to improve its reliability, scalability, and reproducibility.

## Introduction

Genetic screening is a powerful technique for systematically analyzing gene function, reconstructing gene regulatory networks (GRNs), and understanding biological mechanisms^1,2^. These screens are typically conducted in either an individual (“arrayed”) format, where each genetic perturbation is introduced and analyzed separately, or a pooled format, where all perturbations occur simultaneously within a population^2–5^. While both approaches have distinct advantages, recent advances in pooled screening have greatly enhanced the ability to evaluate gene function at a genome-wide scale in mammalian cells^1–3,6–11^. Arrayed screens allow for more detailed molecular readouts, such as RNA sequencing, but at the cost of lower throughput and a highly labor-intensive process, as individual gRNAs must be physically separated^12–14^. In contrast, pooled screening are more efficient and scalable and overcomes these shortcomings of arrayed screens, and had been instrumental^5,15,16^. However, a key limitation of pooled screening is its reliance on relatively simple phenotypic readouts, such as cell viability or the expression of a few exogenous reporters, which provide population-averaged measurements rather than cell-specific insights^17–19^. Additionally, pooled screens are not well suited for complex molecular analyses like transcriptome profiling, which is one of the most comprehensive measures of cellular response^20,21^. Balancing the strengths of both screening approaches is crucial for refining genetic screening methodologies and advancing our understanding of gene function in complex biological systems.

In recent years, with the advancements of CRISPR techniques and single-cell level quantification approaches, pooled CRISPR screening has emerged as a powerful and widely used method, addressing the shortcomings of the traditional pooled screening approach. These approaches allow detecting each cell’s perturbation status along with the corresponding single-cell phenotype at different omics layers^2,3,6,7^. The initial pooled CRISPR screenings approach combines CRISPR-based perturbation to gene expression signatures for individual gene perturbation in a complex pool of cells^2,3,7^. These revolutionary experiments have made significant advances in comprehending the functions of hundreds of genes simultaneously, and helped in uncovering their cellular impact, deciphering gene regulatory networks, and shedding light on several biological mechanisms. Recent developments are now being enhanced to encompass all major omics techniques, including chromatin accessibility, protein concentrations, and imaging approaches that provide novel insights into cell phenotypes^22–27^.

Although the initial approaches were based on CRISPR knockout (CRISPR KO) techniques, recent advancements in CRISPR approaches offer a range of options, including gene knockdown (CRISPR interference, CRISPRi) and activation (CRISPRa) of the genes of interest^28–30^.

CRISPR KO generates permanent gene disruption by introducing small insertions or deletions (indels) that cause a frameshift mutation, and is widely used in genome-wide screens to identify essential genes and studying loss-of-function mutations. In contrast, CRISPRi reduces but does not eliminate gene expression, making it ideal for studying essential genes where a complete knockout would be lethal and for reversible gene modulation. In addition, in CRISPRi is advantageous in avoiding genetic variability during indel formation and non-specific toxicity due to DNA cutting^31,32^. However, despite this advantage, CRISPRi has also its own shortcomings as it requires continuous expression of dCas9 (dead Cas9) and guide RNAs to maintain gene inactivation. These CRISPR-based methods are now being integrated with single-cell approaches to capture cell states influenced by specific perturbations^33–35^. CRISPRi has been successfully applied in combination with single-cell omics approaches. The first pooled CRISPRi perturbation, perturb-seq^2^, used barcodes to identify cells expressing specific gRNAs. Subsequently, other approaches emerged that utilized sgRNA capture and used them as barcodes (CROP-seq)^3^. Despite this development, similar to other pooled perturbations approaches, the field is still emerging. Several improvements and standardizations from experimental to analytical points are still needed to be implemented.

Here, we evaluated the efficiency of CRISPRi-CROP-seq by targeting a focused set of transcription factors (TFs) and generating high-coverage single-cell transcriptomic data in K562 cells. Using this approach, we perturbed six TFs—LMO2, TCF3, LDB1, MYB, GATA2, and RUNX1—and achieved efficient and reproducible knockdown for only three TFs(MYB, GATA2, and LMO2). This enabled us to map their transcriptional targets and reconstruct gene regulatory networks (GRNs), uncovering both known and novel regulatory relationships, with key regulons independently validated. To assess the generalizability of our approach, we analyzed publicly available pooled CRISPRi single-cell screen datasets and found that a substantial fraction of sgRNAs failed to induce effective knockdown, highlighting the variability in perturbation efficiency.

Our findings demonstrate that CRISPRi-CROP-seq is a promising strategy for dissecting gene function and regulatory circuitry at single-cell resolution. However, improvements in perturbation consistency, sgRNA design, and experimental optimization are still necessary to enhance its robustness and scalability for broader applications in functional genomics.

## Results

### Pooled CRISPRi screen with direct 5’ CRISPR capture sequencing

In order to assess the effectiveness of pooled CRISPR-based gene inactivation (CRISPRi) and capture the subsequent impact on phenotype at the single-cell transcriptome level, we used a modified version of CROP-seq^28,36^. This modified CROP-seq (CRISPRi-CROP-seq) contains a scaffold sequence (sgRNA^E+F^, which combines an A-U flip with a 5 bp hairpin extension) that can improve the knockdown (KD) efficiency and can be directly captured using the 10X genomics “Single Cell 5’ CRISPR Screening Assay” (Figure 1A). In the final cDNA library of this assay, two distinct cDNA libraries are simultaneously produced: the mRNA library and the sgRNA library. This direct capture approach is suggested to enhance sgRNA capture sensitivity, as it enables direct capture of both polymerase II and III transcripts, thereby enhancing the accuracy of the results. For this experiment, we selected 7 transcription factors (TFs) known to play key role in the pathogenesis of acute myeloid leukemia (AML) based on: the analysis of a patient cohort^37^, matching with TCGA data^38^, and a thorough literature search. These TFs also show high expression levels in AML patients. The selected TF are LMO2, TCF3, LDB1, MYB, GATA2, and RUNX1 which are all expressed in K562 cells. Additionally, FLI1, is included as a negative control, since it is not expressed in K562 cells. To conduct a comprehensive and thorough investigation of the CRISPRi-CROP-seq, we decided to limit the number of genes in the pool to these seven genes and generate a detailed output.

**Figure 1.**
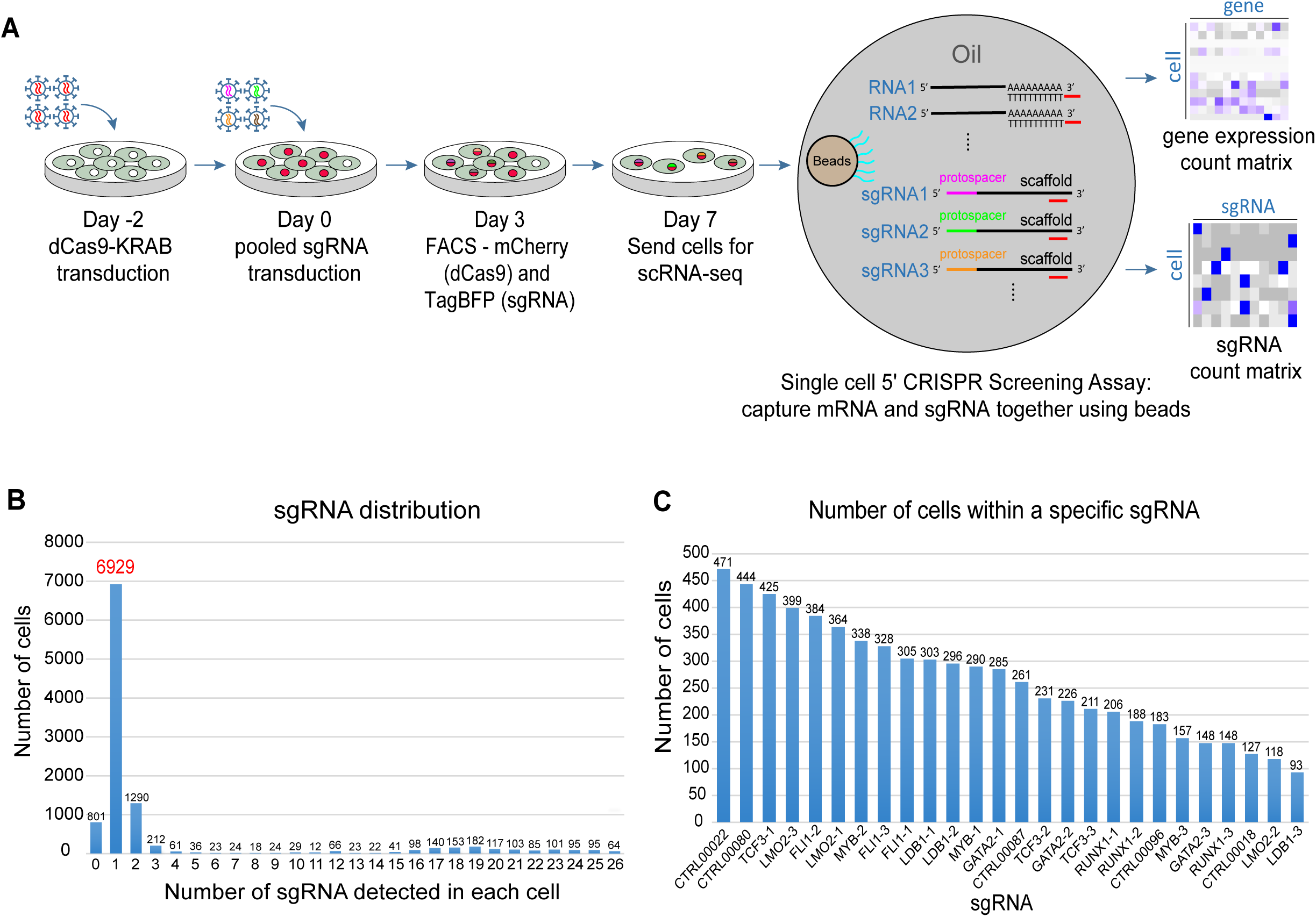
Overview and result of direct 5’ CRISPR capture sequencing. (A) Workflow of the pooled CRISPRi screen with Single Cell 5’ CRISPR sequencing in k562 cells. (B) Distribution of the total number of sgRNAs detected per cell. (C) Number of cells assigned to each sgRNA.

Aiming to inactivate these genes in a pool, we selected 26 sgRNAs: 3 gRNA per gene from the Dolcetto library^39^, and 5 non-targeting sgRNAs from literature^3^. After cloning the sgRNA pool into a lentiviral vector, we confirmed the presence of each gRNA by sequencing the lentiviral vector plasmid library (Supplementary Figure 1A). Subsequently, the sgRNAs were packaged into lentiviruses, transduced into K562 cells, and the presence of each gRNA was confirmed by sequencing amplified sgRNA sequences extracted from the genomic DNA (Supplementary Figure 1B). Next, we transduced these pooled sgRNA, after packaging lentivirus vector, into K562 cells at day 0 (D0), which had been transduced with dCas9 -KRAB-ZIM3 lentivirus two days earlier (D-2). After Fluorescence-activated cell sorting (FACS), sorting double-positive cells at D3, using TagBFP (for cells with sgRNA) and mCherry (for dCas9-KRAB), we performed single cell sequencing at D7 using the 10X genomics platform (Figure 1A).

After sequencing, we recovered 10848 cells for sequencing with two count matrices: the mRNA matrix and the sgRNA matrix. For the gene expression count matrix, we generated a total of 417,3 M reads, with an average of 38,477 reads per cell and a median of 4,623 genes per cell. For the sgRNA matrix (capturing sgRNA), we generated 94,9 M reads, and around 85% of them were usable (reads with a recognized protospacer sequence; a valid unique molecular identifier (UMI); and a cell-associated barcode). The median sgRNA UMIs per cell is 3535. After assigning sgRNA to cells, we found 6929 cells (63.9%) harboring only one sgRNA (Figure 1B), 1290 cells (11.9 %) carrying two sgRNA, and 212 cells (2%) with three sgRNA. Additionally, we also observed a relatively few numbers of cells carrying more than 3 sgRNA, and 801 (7.4%) cells without any sgRNA (with protospacer sequence identified as background). When we assessed the number of cells that are associated with a specific sgRNA, we observed that the highest number was 471, while the minimum count was 93 (Figure 1C), which is sufficient for subsequent analyses. Furthermore, these results demonstrate that the “CRISPR Screening Assay” enables the capture and sequencing of sgRNA with high quality and efficiency.

To conclude, the modified CROP-seq (CRISPRi-CROP-seq) enables the cloning and transduction of gRNAs in a pool. Moreover, it allows direct capture of sgRNA from the cDNA library without the need for a separate sgRNA enrichment step. Furthermore, in the cell line we tested (K562), after sequencing, we recovered a significantly high number of cells with a single sgRNA, which is crucial for subsequent analysis steps.

### Efficiency of gene inactivation in single cell experiments is not efficient

After confirming the ability of the modified CRISPRi-CROP-seq to successfully deliver one sgRNA per cell, in majority of cells, and capture the transcriptome profile in single-cells together with the sgRNA, we next evaluated the efficiency of gene inactivation. To start the analysis, we first determined the expression level of the gRNA in the cells where it was identified -to ensure that the sgRNA is expressed in the cells where it was delivered and captured after single-cell sequencing. For all the captured sgRNA, we observed the expression of the sgRNAs with a median UMI count of 3535 (Figure 2A). In this analysis, we also noticed that even for the same gene, the number of captured sgRNAs per cell varied among the three different sgRNAs, and similar variability was also evident among sgRNAs targeting different genes (Figure 2A).

**Figure 2.**
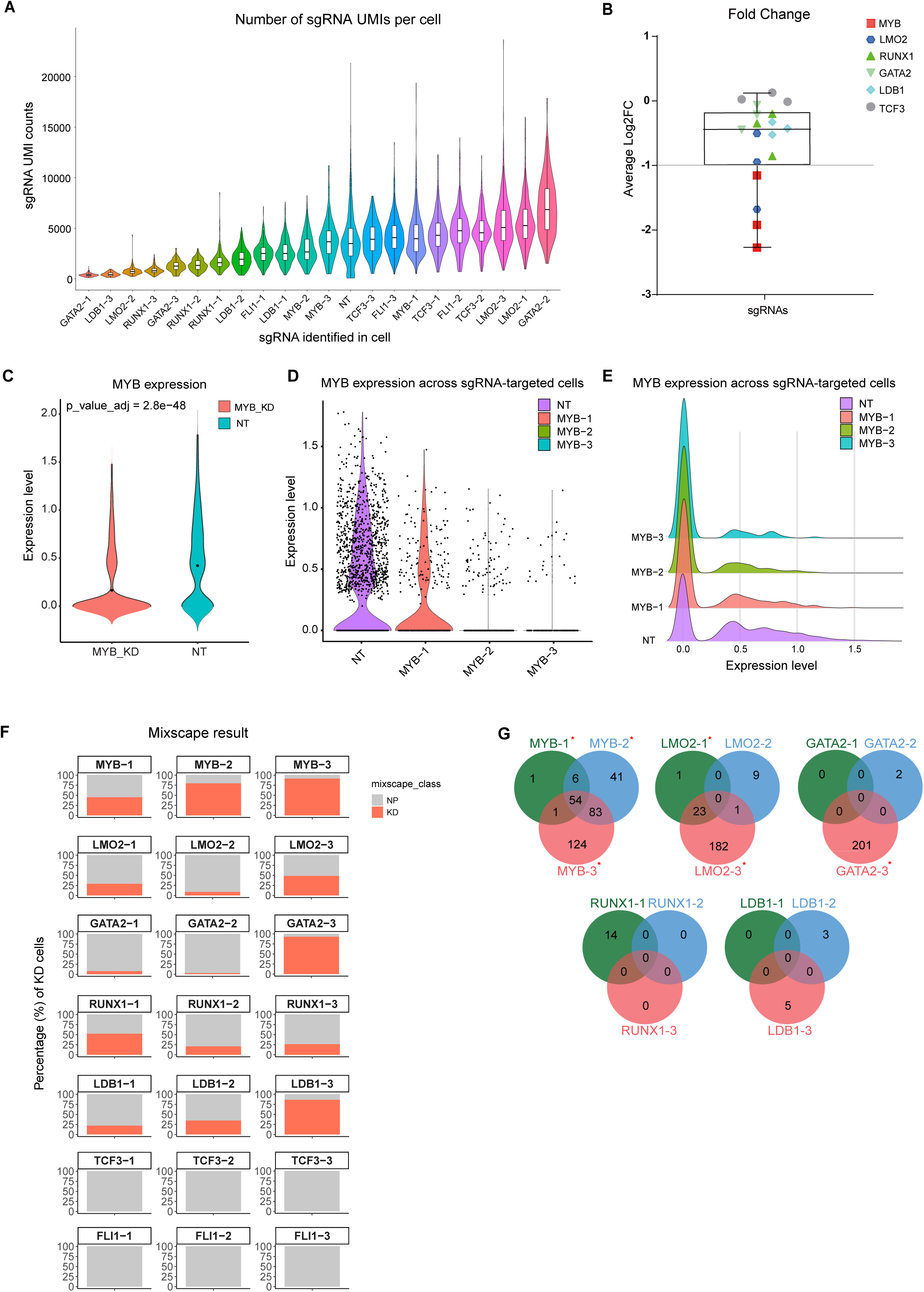
Efficiency of gene inactivation. (A) Violin plot showing UMI counts per cell for each sgRNA. NT: non-targeting control cells. (B) Fold change in expression for the 18 sgRNAs used in the experiment. (C-E) Violin plot showing MYB expression across cells transduced with different MYB-targeting sgRNAs. (F) Percentage of knockdown (KD) cells for each sgRNA as estimated by Mixscape. (G) Overlap of DEGs identified from the three separate sgRNAs targeting each gene.

To understand the KD (knockdown) efficiency on the sgRNA level, we calculated their FC (Fold Change) separately. The result (Figure 2B) showed that only 4 sgRNAs (MYB-1, MYB-2, MYB-3, LMO2-3) achieve efficient KD (with log2FC < -1). To further visualize these findings, we generated violin plots at both the sgRNA and gene levels (Figure 2C-E, Supplementary Figure 2). The results revealed considerable variability among sgRNAs targeting the same gene.

To gain a more comprehensive view we then used the Mixscape computational framework that enhances signal-to-noise ratio in single-cell perturbation screens, and enable the separation of a cell mixture into perturbed and non-perturbed cells^26^. We performed Mixscape analysis at sgRNA level, to show the impact of each sgRNA on target gene expression (cells with each sgRNA) and found that 15 out of the 21 sgRNA perturbed (KD) the target gene in varying number of cells (Figure 2F). For some sgRNA that target the same gene, we observed a large variability. For example, for the sgRNAs targeting GATA2, the variability of cells, where the sgRNA targeting GATA2 were captured and the target gene was perturbed, varied between 5% to 90% (Figure 2F). For other genes, such as MYB, this variability is limited (Figure 2F). For other genes, such as TCF3, despite the sgRNAs is expressed (Figure 2A) and also captured, we could not identify cells where the gene is perturbed. For FLI1, that we used as a negative control as it is not expressed in the K562 cells, we were able to capture cells where the sgRNA targeting the gene is present, but as expected did not show perturbation (Figure 2F). This raises a question about the efficiency of the system. Even for genes that have a significant perturbation, when the sgRNAs were pooled, there is a large variability between the efficiency of sgRNAs (Figure 2B, F).

To further understand the contribution of each sgRNA, we tested if each of the 3 separate sgRNAs targeting the same gene produce similar differentially expressed gene (DEG) patterns in cells. To test this, for each individual sgRNA, we performed differential expression analysis by comparing cells showing efficient KD with cells carrying a non-targeting (NT) sgRNA. In these analyses, we observed that different sgRNAs generate different sets of DEGs (Figure 2G). The overlap of DEGs by these sgRNA varies per targeted gene and, in some cases, this overlap is dependent on the efficiency of the sgRNA. For sgRNAs targeting the MYB gene, we observed the highest DEGs overlap between the sgRNAs targeting it (Figure 2G), suggesting that all three MYB sgRNAs are effective. For LMO2, the sgRNA LMO2-3 yielded the highest number of DEGs (Figure 2G). In contrast, the sgRNA LMO2-1 and LMO2-2 have a relatively low number of DEGs. Notably, there is a substantial overlap in DEGs between LMO2-1 and LMO2-3, suggesting that LMO2-1 is relatively efficient. However, LMO2-2 shows minimal overlap with the other two sgRNAs, indicating low efficiency. In the case of GATA2, only one sgRNA (GATA2-3) generated 201 DEGs. The other sgRNAs targeting RUNX1 and LDB1 did not yield significant results, and we only obtained a very low number of DEGs. In general, based on this differential gene expression analysis, only 6 out of the 18 sgRNAs had a significant impact.

To further investigate the impact of these perturbations, we focused on the 3 genes (MYB, GATA2 and LMO2) that show significant perturbation. We extracted all the KD cells from these 6 efficient KD sgRNAs and combine them on the gene level. We first clustered the cell population and plotted them using Uniform Manifold Approximation and Projection (UMAP), together with the cells with control sgRNAs that does not target any gene (NT), and we observed a clear separation of the cell where we perturbed the genes forming distinct clusters (Figure 3A), indicating the impact of the perturbation. We then performed differential expression analysis and plotted these DEGs (cutoff: |log FC| >0.25 adjusted p-value < 0.05 ), and identified DEGs (914 for MYB ,1364 for LMO2, 236 for GATA2, see Supplementary Table 1). The DEGs were visualized using a heatmap and showed a clear clustering of the cells based on the expression of these DEGs (Figure 3B, supplementary Figure 3). Among them, we found MYADM, STAT5A, LMNB1, TMX2, GATA2, and MYC, all of which have been previously reported as MYB target genes^40,41^.

**Figure 3.**
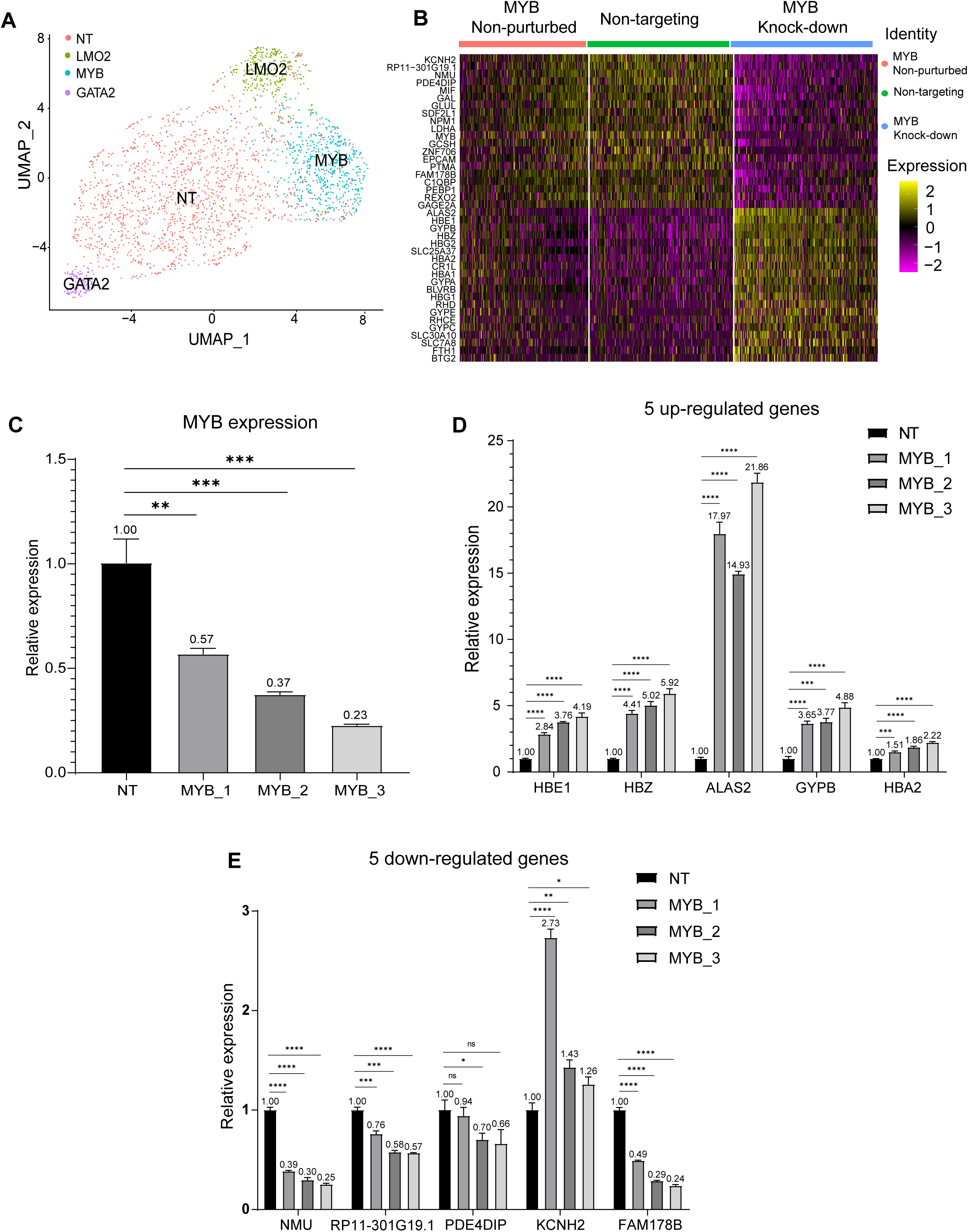
Validation of CRISPRi-CROP-seq perturbation results. (A) UMAP plot showing effective gene knockdown cells across all six sgRNAs and non-targeting (NT) control cells. (B) Heatmap comparing MYB knock-down (KD) and non-perturbed (NP) cells (as estimated by Mixscape) to NT cells. (C) RT-qPCR analysis in K562 cells of MYB expression in MYB KD versus NT cells. (D) RT-qPCR validation of five upregulated DEGs identified in MYB KD cells compared to NT controls. (E) RT-qPCR validation of five downregulated DEGs following MYB knockdown. Data are presented as mean ± SD. Statistical significance was determined using two-tailed Student’s t-test. Significance levels are indicated as follows: ns, not significant; * p < 0.05; ** p < 0.01; *** p < 0.001; **** p < 0.0001.

We next validated these observations using RT-qPCR to confirm the downregulation of the targeted gene by the sgRNA and also the DEGs identified in the single-cell experiment following similar CRISPRi experimental protocol we applied for the single-cell experiment. We perturbed (KD) targeted genes separately by transfecting the sgRNA plasmid into k562 cells carrying dCas9. After sorting cells using Fluorescence-activated Cell Sorting (FACS) on day 2 and RNA extraction on day 3, we performed RT-qPCR to quantify the sgRNA targeted gene expression and DEG expression for each sgRNA. In this validation, we observed the expected downregulation of the target gene as observed in the single cell expression analysis (Figure 3C, Supplementary Figure 4A-B).

To validate our results, for MYB gene, we selected 5 DEGs that were upregulated (HBE1, HBZ, ALAS2, GYPB, HBA2) and 5 DEGs were downregulated (NMU, RP11-301G19.1, PDE4DIP, KCNH2, FAM178B) in the single-cell experiment. MYB is a key TF in erythropoiesis and hematopoietic stem cell regulation and cancer^42^. In this validation experiment, similar to what we observed at the single-cell RNA level, we observed comparable efficiency for each sgRNA targeting MYB (Figure 3C). Additionally, we validated the expression changes in these MYB target DEGs using RT-qPCR, supporting the impact of MYB inactivation observed in the single-cell experiment (Figure 3D, 3E). We performed a similar analysis for GATA2 and LMO2, two key transcription factors involved in hematopoiesis, vascular development, and immune system function^43,44^. For GATA2, we selected the 10 DEGs, with five upregulated genes (ID3, APOE, PYCARD, CD52, and SPI1) and five downregulated genes (PRDX1, SNAP29, PAFAH1B3, HBD, and KRT19) (Supplementary Figure 4C-D). Similarly, for LMO2, we validated the expression of 10 DEGs: five upregulated genes (ID3, CD24, VIM, APOE, and SNCG), and five downregulated genes were HBZ, HBG1, GAL, KLF1, and HMBS (Supplementary Figure 4 E-F). In all instances, we were able to validate the perturbation of the sgRNA target genes (MYB, GATA2 & LMO2) and the DEGs using RT-qPCR.

We next performed gene enrichment analysis using the DEGs, and observed that for genes that are downregulated upon MYB perturbation, we observed enriched terms related to ribonucleoprotein complex biogenesis, metabolism of RNA, translation and protein maturation. For upregulated genes the enriched terms included erythrocyte development, porphyrin-containing compound metabolic process and regulation of hemoglobin biosynthetic process (Supplementary Figure 5A). This is consistent with previous findings on the MYB gene, which show that high MYB expression can inhibit cell differentiation in erythroid cell lines^42^ and is essential for malignant self-renewal^45^. To further validate our findings, we analyzed a MYB ChIP-seq dataset performed on K562 cells^40^. The analysis revealed clear MYB occupancy around transcription start sites (TSS, ±5kb) of down-regulated DEGs, whereas no significant enrichment was observed around the TSS (±5kb) of up-regulated DEGs (Supplementary Figure 5B). This is also consistent with the established role of MYB as a transcriptional activator^46^. In a similar fashion, we performed enrichment (Supplementary Figure 6A) and ChIP-seq (Supplementary Figure 6B) anlysis^47^ for GATA2-regulated DEGs. After enrichment analysis, we found that the downregulated GATA2 DEGs were mainly enriched in ribonucleoprotein complex biogenesis, PID MYC activation pathway, chaperone-mediated protein folding and cell population proliferation (Supplementary Figure 6A). For the upregulated genes, they were enriched in cholesterol metabolism, maintenance of location, lysosome and neutrophil degranulation (Supplementary Figure 6A). When we analyzed GATA2 ChIP-seq data (Supplementary Figure 6B) that was generated on K562 cells, and mapped the occupancy around TSS of the up and down regulated genes, we observed clear enrichment for upregulated genes compared to downregulated genes.

In conclusion, we successfully perturbed (KD) three TFs—MYB, GATA2, and LMO2—out of the six TFs we had targeted in K562 cells, captured the impact of these perturbations on their target genes, and verified the results. However, despite this moderate success, these results indicate that, the efficiency of the CRISPRi-CROP-seq experiment depends on several aspects that need several considerations.

### Impact of perturbations on gene regulatory networks (GRNs)

Perturbation of TFs is expected to alter GRNs, and since we observed significant changes in gene expression after knocking down MYB, GATA2, and LMO2, we investigated the impact of these perturbations on GRNs to better understand their effects. We applied SCENIC^48^, a tool that infers GRNs from scRNA-seq data. In this analysis, we identified a total of 118 high-confidence regulons (a group of genes regulated by the same regulatory protein). We first analyzed regulon activity (regulon activity score, RAC) at both the sgRNA level (Supplementary Figure 7A) and gene level (Figure 4A). At the sgRNA level, most efficient sgRNAs exhibited distinct high or low regulon activity, except for MYB-1, which showed a similar but weaker activity pattern compared to MYB-2 and MYB-3. This result aligns with Fold Change and Mixscape analyses (Figure 2B and F), and confirming MYB-1 is less effective than MYB-2 and MYB-3. At the gene level (grouping sgRNA targeting the same gene), KD cell groups displayed distinct high-or low-activity regulons. As expected, MYB, LMO2, and GATA2 regulons showed reduced activity in their respective MYB, LMO2, and GATA2 KD groups, validating the specificity and efficiency of the knockdowns. To quantitatively assess how specifically each regulon is active in particular cell clusters, we performed regulon specificity (shown as regulon-specific score, RSS) analysis across all clusters (Figure 4B). The results revealed that each KD cluster had distinct high-specificity and high-activity regulons, as indicated by high Z-scores. For example, the STAT6 regulon exhibited a significantly higher Z-score in the GATA2 KD cluster compared to other clusters, suggesting a strong activation following GATA2 KD, a pattern not observed in MYB KD or LMO2 KD.

**Figure 4.**
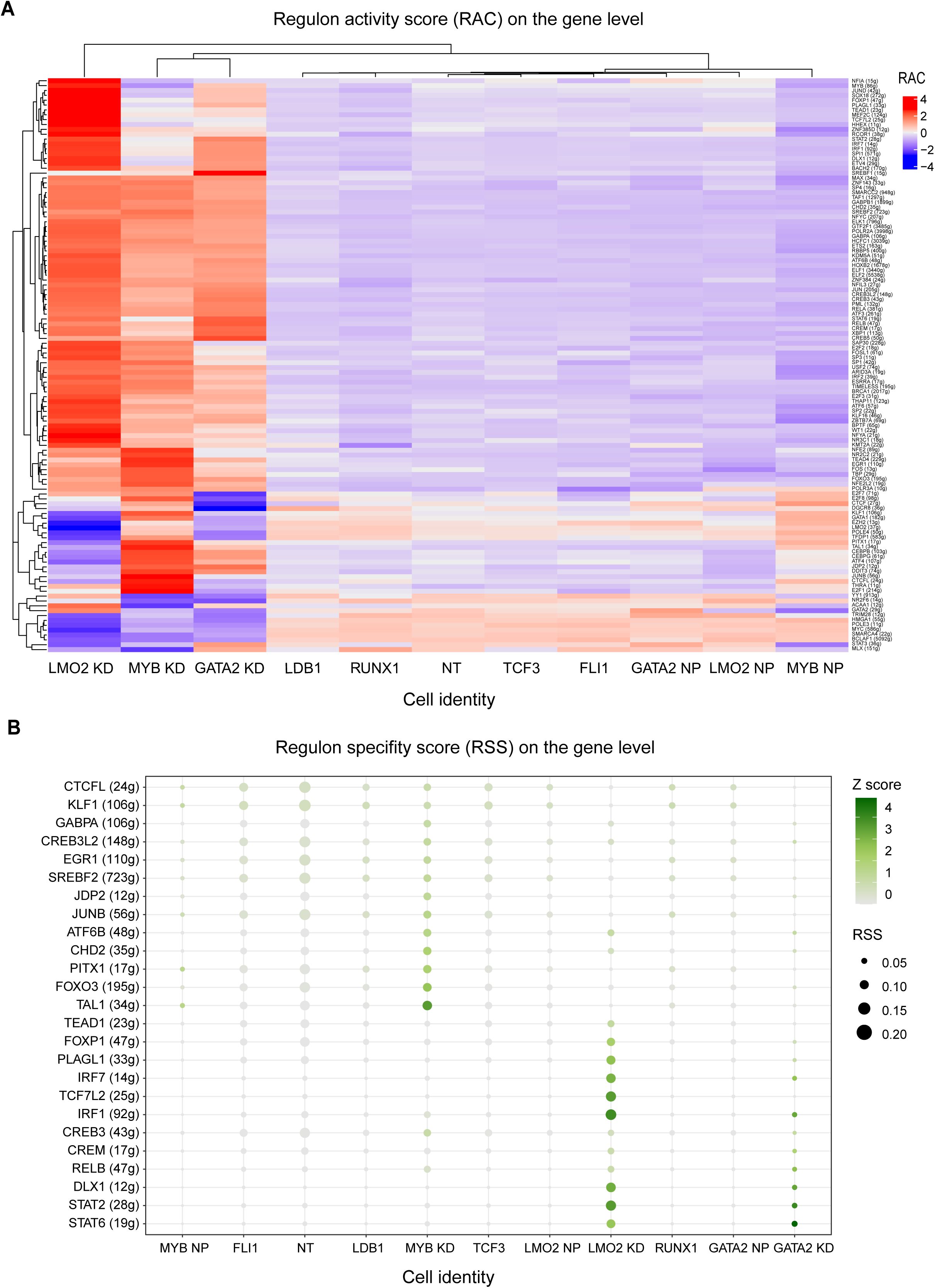
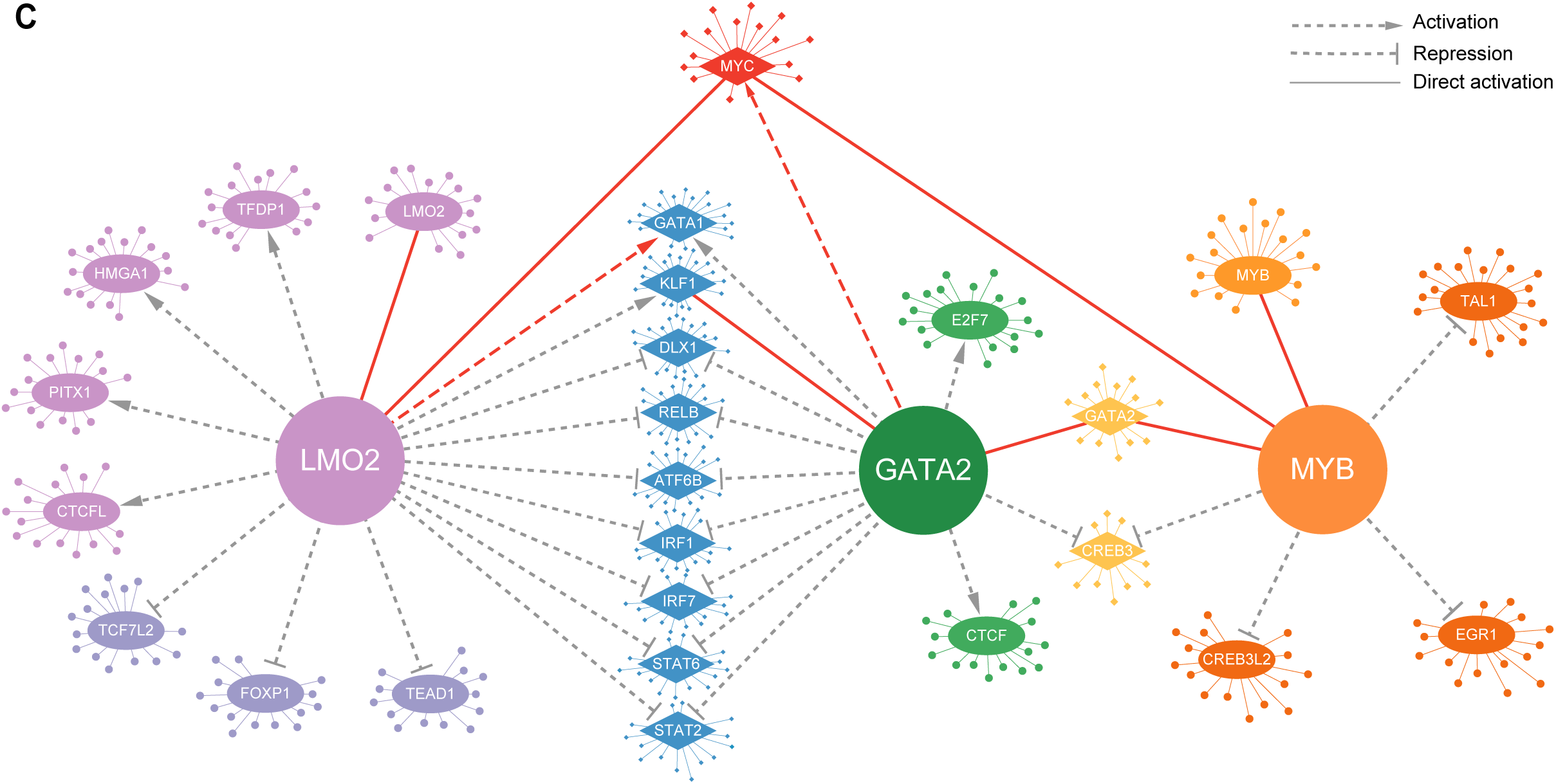
Gene regulatory networks. (A) Regulon activity at perturbed gene (gene level summary, an average of 3 sgRNAs’ KD cells). The X-axis represents cell identity (based on sgRNA that is identified), while the Y-axis represents regulons. “NT” refers to a combination of five non-targeting sgRNAs. (B) Regulon-specific score at the gene level. A higher Z-score indicates greater specificity of that regulon for the given cell group. (C) Gene Regulatory Networks based on this current sgRNA screen, and solely focused on LMO2, GATA2 and MYB. Grey dotted lines indicate putative regulatory connections (activation or repression). The red dotted line represents a previously reported indirect connection. Solid lines indicate direct regulatory interactions; among them, red solid lines correspond to interactions supported by published evidence. Except for the central genes LMO2, GATA2, and MYB, all other genes are regulons. For example, TAL1 is repressed by MYB (either directly or indirectly) and directly activates many genes (represented by the small dots connected to TAL1). Note that the number of small dots is symbolic and does not reflect the actual number of target genes.

Next, for each cluster (MYB KD, GATA2 KD, and LMO2 KD), we screened important regulons based on two criteria: 1) low-activity regulons (RAC < -1); (2) high-specificity and high-activity regulons (RSS Z-score > 0.5). This initial filtering yielded 39 regulons for GATA2 KD, 40 for LMO2 KD, and 42 for MYB KD (Supplementary Table 2). To further refine the list of relevant regulons, we calculated the fold change (FC) of the core transcription factors (TFs) in each regulon (e.g., MYC in the MYC regulon), as these TFs are expected to show corresponding changes in expression. Regulons were retained only if their TF exhibited a significant expression change, defined as an absolute log₂ fold change (|log₂FC|) greater than 0.5. This finally, resulted in 23 key regulons, and by leveraging the connections among these regulons, we constructed a gene regulatory network (GRN) (Figure 4C). In this network, the most strongest connection was between the LMO2 and GATA2, which is consistent with previous findings that identified LMO2 as interactor of GATA2^49^. This network analysis not only confirmed several well-established direct gene interactions (e.g., MYB – MYC^40,50^, MYB – GATA2^46^, GATA2 – KLF1^51^, and LMO2 – MYC^52^), but also predicted indirect interactions, such as GATA2 – MYC^53^ and LMO2 – GATA1^54^. For the GATA2 – MYC interaction, one study demonstrated that pathway analysis of RNA-seq data following GATA2 deletion in AML leukemic stem cells (LSCs) revealed downregulation of MYC target genes. This finding suggests that GATA2 may function as a positive regulator of the MYC pathway, rather than directly regulating MYC itself. Regarding the LMO2 – GATA1 interaction, it has been shown that LMO2 contributes to the expression of GATA1 target genes by modulating the assembly of the GATA1–SCL/TAL1 transcriptional complex at endogenous loci. The presence of these known interactions further reinforces the accuracy and biological significance of our GRNs.

To further validate these inferred GRNs, we performed independent knockdown experiments for LMO2, and GATA2 in K562 cells. We then selected a subset of genes (based on the observed expression change in the GRN analysis) for RT-qPCR analysis to assess their expression changes. The results demonstrated that the majority of these genes exhibited the expected upregulation or downregulation (Supplementary Figure 7B-C). For example, LMO2 represses TFs such as IRF1, IRF7, and FOXP1, thus genes within these regulons are expected to be upregulated upon LMO2 KD. Consistent with this, we observed increased expression of ANK3 (IRF7 regulon), FN1 (IRF1), and IGF1 (FOXP1) in LMO2-repressed K562 cells. Similarly, GATA2 KD resulted in upregulation of AIF1 (STAT2 regulon), CD53 and PYCARD (IRF1), CORO1A (STAT6), and SPI1/TPM4 (CREB3), consistent with its repression role. In contrast, KRT19 (GATA1 regulon) was downregulated, as expected from GATA2’s activation of GATA1. These results provide strong support for the reliability of our inferred GRNs.

In conclusion, we constructed GRNs for MYB, GATA2, and LMO2 using gene perturbation and identified key regulons and target genes regulated by these transcription factors. We were also able to validate these observations independently. Our results reveal both novel interactions and previously established ones, highlighting the power of this approach.

### Pooled CRISPRi based gene perturbation and single-cell readout datasets have low efficiency

In recent years, several pooled CRISPRi single-cell screen experiments have been published and utilized to address various questions^55–57^. In the CRISPRi-CROP-seq experiment that we performed to assess the efficiency of this approach and unravel the regulatory network of the genes we targeted, the low efficiency we observed in our settings prompted us to question the reliability of other pooled CRISPRi single-cell screen experiments that have been recently published. To investigate and compare our results, we acquired two publicly available, published datasets^34,58^ that conducted pooled CRISPRi screen with single-cell sequencing experiments with direct capture sequencing, which is similar to our approach. The first dataset^34^ (Dataset one, Wu et al. 2022, conducted a pooled CRISPRi single-cell screen experiment in induced pluripotent stem cells (iPSCs) using 480 single-guide RNAs (sgRNAs) to target and perturb 120 coding genes and 120 long non-coding RNAs (lncRNAs). Since our experiment focused primarily on coding genes, we retained the cells with sgRNAs targeting only coding genes and excluded those targeting lncRNAs. The second dataset^58^ (Dataset two, by Joseph M.R et al. 2022), performed pooled CRISPRi single-cell screen in K562 cells using 384 sgRNAs targeting 128 genes, aiming to improve the CRISPRi efficiency by dual-sgRNA design.

We analyzed these datasets using a similar pipeline and approach used to analyze our CRISPRi-CROP-seq experiments data. First, we quantified the number of sgRNA that were used in the pool and what was obtained after the single cell sequencing. For dataset one, <u>all</u> the 240 sgRNA were captured, that is similar to what was published by the authors (Figure 5A). For dataset two, 384 sgRNAs, 377 were captured, also as initially reported by the authors (Figure 5A). Next, we analyzed the distribution of sgRNA per cell. For dataset one (Figure 5B), we obtained 78929 cells with one sgRNA, 27063 cells with 2 sgRNA and 8239 cells with 3 sgRNA. For dataset two (Figure 5C), we obtained 57989 cells with one sgRNA, 11799 cells with 2 sgRNA and 2571 cells with 3 sgRNA. This distribution of sgRNA is also in agreement with what the authors reported. Additionally, we also counted the median UMI for each sgRNA (Figure 5D-E), ranges from 1 to 600 in Dataset One and from 1 to 1200 in Dataset Two. This variation shows the variability between sgRNAs.

**Figure 5.**
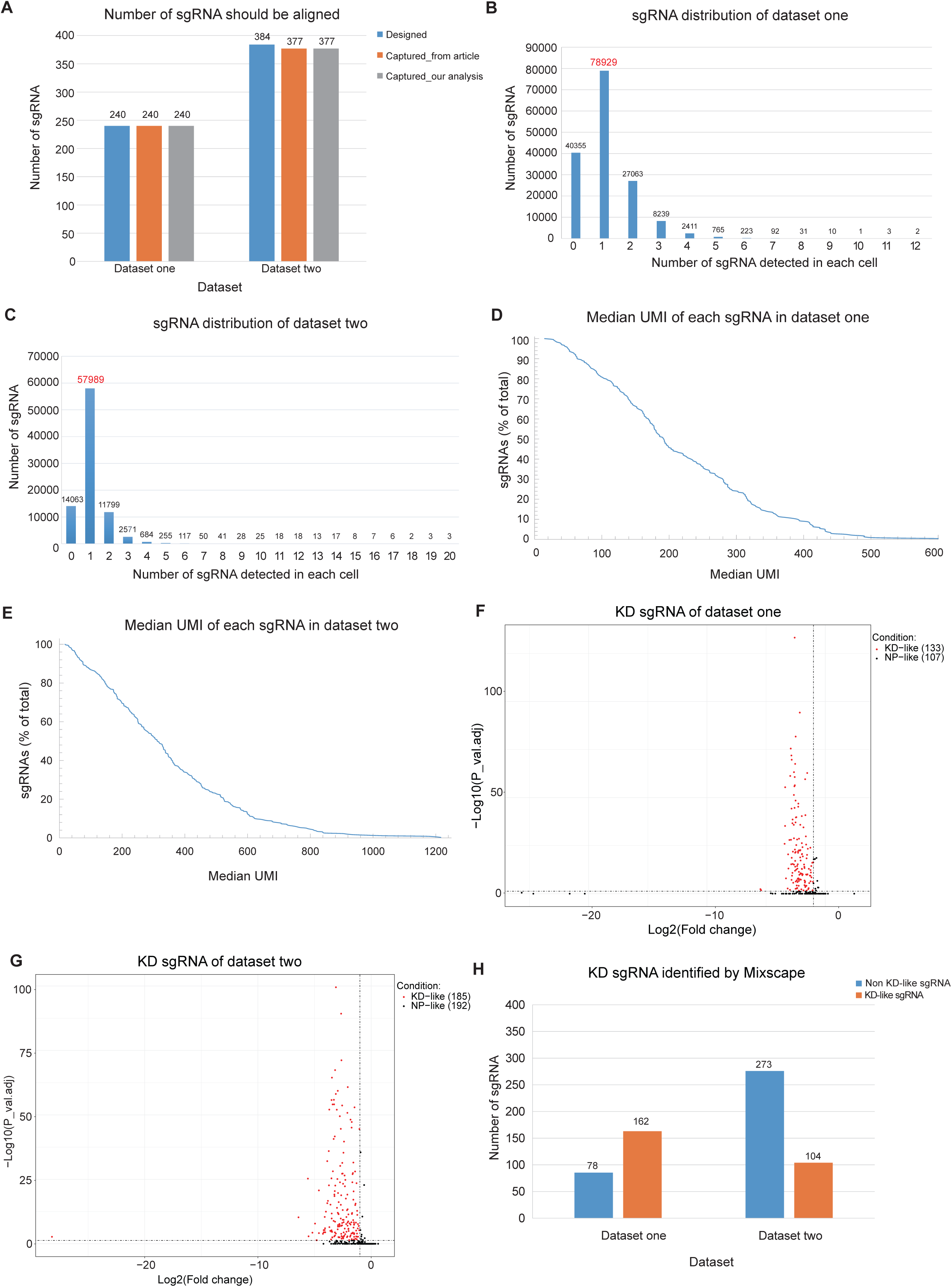
Analysis of two published Perturb-seq CRISPRi datasets. (A) Number of sgRNA captured in the different analysis. (B,C) Distribution of the number of sgRNA detected in each cell for the two datasets. (D,E) Median UMI of each sgRNA for the two datasets. (F,G) Volcano plots which represent the number of sgRNA surpassing the Fold Change and p_val_adj thresholds in the two datasets. (H) Bar plots which represent the number of efficient KD sgRNA identified by the Mixscape program.

To estimate the efficiency of gene inactivation by sgRNA, we first calculated FC and adjusted P-value for target gene of sgRNA (log2FC < -1 and adjusted P-value <0.05). Only sgRNA that passes the threshold can be identified as KD-like. For dataset one (Figure 5F), in iPSC, from the 240 sgRNA, only 133 show efficient gene inactivation, and for dataset two (Figure 5G), in K562 cells, only 185 out of the 377 sgRNA show significant gene inactivation. This indicates, in both datasets, only around half of the sgRNAs used are able to inactivate the target gene. Next, we used the Mixscape framework to determine the target gene perturbation efficiency at sgRNA level. For dataset one (Figure 5H, Supplementary Figure 8A), 162 sgRNAs were found to have efficient gene perturbation, and 104 sgRNAs for the dataset two (Figure 5H, Supplementary Figure 8B). When we plotted the impact of these sgRNA using UMAP (Supplementary Figure 8C-D), to visualize their impact and we observed that only few genes having a significant impact and differ from the sgRNA in the pool. Although this visualization is not quantitative, it demonstrates that only a few genes have a significant impact, which further confirms the low efficiency of the experiments. A large proportion of sgRNAs do not have a significant impact on gene perturbation.

In the next step, we conducted a more stringent analysis by integrating the fold change-based analysis of the target gene with the results from the Mixscape analysis. Only sgRNAs identified by both methods were classified as KD sgRNAs (Supplementary Figure 8E-F). For dataset one, in the iPSC dataset, only 97 out of 240 sgRNAs KD efficiently. For the K562 dataset, only 56 out of 377 sgRNAs KD efficiently. This again points to the fact that the limited efficiency of CRISPRi-CROP-seq experiments.

To conclude, in this analysis we show that, in these published datasets a significant proportion of the sgRNA does not produce significant perturbation to inactivate the targeted gene. This is in line with efficiency problems we observed in CRISPRi based single cell pooled screens.

## Discussion

Massively parallel gene perturbation techniques have become indispensable for elucidating gene function, mapping gene regulatory networks (GRNs), and uncovering intricate molecular pathways. In our study, we employed CRISPRi-CROP-seq to interrogate the roles of six transcription factors (TFs), yielding detailed insights on three of them. By individually inactivating each TF, we identified their downstream target genes, reconstructed GRNs, and delineated the interrelationships among these regulators. Furthermore, we evaluated the efficiency of this powerful not yet standardized approach. Our results demonstrate that while CRISPRi-CROP-seq holds considerable promise, significant optimization and rigorous benchmarking, including a reevaluation of existing studies, are essential to fully harness its potential.

To investigate the function and regulatory relationships of six TFs—MYB, GATA2, LMO2, RUNX1, LDB1, and TCF3—that we selected based on the high expression of these genes in AML patients. These factors play crucial roles in erythropoiesis, hematopoietic stem cell regulation, and cancer^52,59–63^. While previous studies have examined these TFs in isolation, our study represents the first attempt to concurrently dissect their functions and interrelationships within a single experimental framework. This unified approach minimizes confounding variables by allowing us to assess multiple gene perturbations under identical conditions. As a result, we gained insights into the roles of three TFs—MYB, GATA2, and LMO2— demonstrating that their downstream target genes are integrally involved. For instance, MYB-associated differentially expressed genes (DEGs) were linked to biological processes consistent with its known function, and these findings were independently validated using RT-qPCR. Moreover, integrating MYB ChIP-seq data revealed significant enrichment of MYB binding at the transcription start sites of genes down-regulated in the MYB knock-down condition in our CRISPRi perturbation, further supporting its regulatory impact. In addition to identifying DEGs, we reconstructed the gene regulatory networks (GRNs) for MYB, GATA2, and LMO2 using CRISPRi perturbation data, and validated these networks by RT-qPCR. This network analysis not only confirmed known regulons but also uncovered novel regulatory circuits that illustrate the intricate interactions among these TFs (Figure 4). One example is the predicted interaction between GATA2 and IRF1^64^. Although no studies have directly reported this connection, one study, in acute promyelocytic leukemia (APL), showed MYC represses IRF1 expression through recruitment of the PML-RARα to the IRF1 promoter. Given that GATA2 may act as a positive regulator of the MYC pathway, it is reasonable to propose that GATA2 could also contribute to the repression of IRF1 expression, potentially through an indirect mechanism involving MYC. However, we also observed one potentially spurious association within the reconstructed GRNs, suggesting that GATA2 may promote the expression of GATA1^65^. This finding appears inconsistent with the well-known “GATA switch” during hematopoietic differentiation. In erythroid lineage, increased GATA1 expression would suppress GATA2, thereby facilitating differentiation^66^. Also, sustained high expression of GATA2 is known to promote leukemia cell self-renew and inhibit differentiation, suggesting that GATA2 should repress, rather than activate, GATA1 expression^67^. The observed positive regulation may therefore be a false positive, possibly resulting from the high sensitivity of the SCENIC algorithm, which can sometimes identify indirect or spurious regulatory links. This highlights the need for cautious interpretation of computationally inferred networks and further experimental validation. Despite this, the other findings of our study—confirming known interactions and identifying novel ones—lay the foundation for future hypothesis generation and investigation.

Although our investigation yielded key insights for three of the six targeted transcription factors, several limitations warrant discussion. First, inherent to single-cell approaches is the incomplete capture of the transcriptome. For instance, our median detection of 4,623 genes per cell is typical for the 10X Genomics platform; however, it raises important questions about whether this value accurately reflects the average gene expression per cell or if low-abundance transcripts are systematically underrepresented. A second, and more significant, limitation concerns the efficiency of the pooled CRISPRi perturbation method, as well as the broader challenges associated with massively parallel gene perturbation. Out of the six genes perturbed, meaningful insights were obtained for only three—an approximate 50% success rate. The low perturbation efficiency observed in our study echoes findings from other published works using similar setups. In our reanalysis of two of these datasets, we noted significantly lower efficiency rates (efficient KD was observed in 97 of 240 sgRNAs in dataset 1 and 56 of 377 sgRNAs in dataset 2) than originally reported, which prompts a careful reevaluation of these approaches. In our experiments, the result is particularly notable given that the experimental design was optimized for a high cell count per sgRNA and deep sequencing coverage. In our work flow, we enriched for cells expressing dCas9 and sgRNA via FACS, employed a low multiplicity of infection to deliver a single sgRNA per cell, and effectively captured the sgRNAs through a 5ʹ capture system (see Figure 1). Despite these measures, visible perturbation was limited to half of the targeted genes. Since every procedural step—from sgRNA delivery to capture— performed as expected, we suspect this limited efficacy is tied to factors intrinsic to sgRNA design. These factors include the sgRNA’s ability to bind target promoters (which may be influenced by the local chromatin state) and its capacity to recruit the necessary repressive complexes. The sgRNAs used in our study were obtained from an existing library that might not fully account for these parameters. Future iterations of CRISPRi-CROP-seq could benefit from, carful design of sgRNA taking local chromatin state in to context, incorporating strategies that allow delivery of multiple sgRNAs per cell (such as delivering two or more sgRNAs in a single construct). Based on our observations, we advocate for a comprehensive benchmarking of all massively parallel gene perturbation methods to validate their reproducibility and enhance overall effectiveness.

In conclusion, our study employed massively parallel gene perturbation via CRISPRi-CROP-seq to investigate six transcription factors in K562 cells, and generated valuable insights into three key TFs— MYB, GATA2, and LMO2—and their interconnected GRNs. By integrating perturbation data with independent validations such as RT-qPCR and ChIP-seq, we confirmed the roles of these TFs. At the same time, our work highlights critical limitations of the current approach. Specifically, challenges with incomplete transcript capture at the single-cell level and the reduced efficiency of sgRNA-mediated perturbation underscore the need for careful optimization in sgRNA design and experimental strategy. We recommend that future studies consider incorporating multiple sgRNAs per cell and implementing rigorous benchmarking protocols to enhance data reproducibility and reliability. Ultimately, our findings not only advance our understanding of TF function in blood cell development but also pave the way for refining massively parallel gene perturbation methods for broader genomic applications.

## Material and Methods

### Cell line production and maintenance

K562 cells were maintained in RPMI160 medium with GlutaMAX (Thermo scientific, catalog no. 61870044) supplemented with 10% fetal bovine serum (FBS), 100 units/mL penicillin and 100 μg/mL streptomycin. HEK239T cells were grown in Dulbecco’s modified eagle medium (DMEM) in 10% FBS, 100 units/mL penicillin and 100 μg/mL streptomycin.

To construct the K562 cell line that can stably express dCas9-KRAB, K562 cells were transduced with lentivirus containing pHR-UCOE-SFFV-dCas9-mCherry-ZIM3-KRAB (Addgene, plasmid #154473) at a multiplicity of infection (MOI) approximately 0.5. Three days post-transduction, mCherry-positive polyclonal cells were isolated through fluorescence-activated cell sorting (FACS, BD FACS AriaII, Supplementary Figure 9).

### sgRNA library cloning and quality control

For each target gene, three sgRNAs from the set A pool in the Human CRISPR Inhibition Pooled Library (Dolcetto)^39^ were chosen. To enhance efficiency, a “G” was introduced at the 5’ of each sgRNA. Additionally, five non-targeting sgRNAs sourced from literature were incorporated into our experiment. All these sgRNAs were synthesized by IDT as 73/72 base single-stranded oligonucleotides structured as follows: GGAGAACCACCTTGTTG (5’ overhang to the mU6 promoter) + G + sgRNA + GTTTAAGAGCTAAGCTGGAAACAGCATAGCAAGTT (3’ overhang to the gRNA scaffold), and ordered as a pool (Supplementary Table 3).

The vector backbone was prepared by digesting the pBA950 plasmid (Addgene, plasmid #122239) with BlpI (NEB, catalog no. R0585S) and BstXI (NEB, catalog no. R0113S) in NEBuffer r2.1 at 37°C for 1 hour. After subjecting the digested vector to gel electrophoresis, it was excised and purified using the gel clean-up kit (Takara, catalog no. 740609.250). The sgRNA library was then constructed following the CROP-seq protocol^3^. Library coverage was assessed by enumerating bacterial colonies on the 1:1,000 dilution plate, revealing that the sgRNA library was cloned at 770x coverage.

The sgRNA sequence was amplified from either the plasmid library or the genomic DNA (gDNA) extracted from cells transduced with the sgRNA library lentivirus. The sequencing libraries were prepared using the CRISPR library preparation method^3^. The libraries were sequenced on an Illumina NextSeq 2000 platform, generating 50-base paired-end reads: Read 1 encompassed the end of mU6 promoter (22 bases), the gRNA spacer (20/21 bases), and the beginning of scaffold (8 bases), while Read 2 only covered the scaffold. sgRNA sequences were aligned to the Read 1 files with a perfect match (no mismatch) to investigate their distribution.

### Lentivirus production

One day before transfection, 7 million HEK239T cells were seeded onto a 10 cm dish in 12 ml Opti-MEM I (Gibco, catalog no. 51985-034) supplemented with 5% FCS and 200 μM sodium pyruvate. Following overnight incubation to attain 90% confluency, HEK 293T cells were transfected using Lipofectamine 3000 reagent (Invitrogen, catalog no. L3000015). To make transfection mixture A, 10.2 μg of the modified pBA950 (containing either one sgRNA or the entire library) or dCas9-KRAB (Addgene, plasmid #154473) was combined with 5.4 μg each of the three packaging plasmids: pMDLg/pRRE (Addgene, plasmid #12251), pRSV-Rev (Addgene, plasmid #12253), and pMD2.G (Addgene, plasmid #12259). This mixture was added to 1.5 ml Opti-MEM I, supplemented with 36 μl P3000 enhancer reagent.

Simultaneously, transfection mixture B was prepared by blending 42 μl Lipofectamine 3000 reagent with 1.5 ml Opti-MEM I. Lipid-DNA complexes were generated by combining transfection mixes A and B, followed by a 20 minutes’ incubation at room temperature. Subsequently, 6 ml medium was removed from the HEK293T cells, and 3 ml lipid-DNA complexes was added. After a 6-hour incubation, the medium was replaced with 12 ml fresh medium. Lentivirus supernatants were harvested 48 hours post-transfection and filtered (0.45 μm). Immediately after collection, lentivirus were concentrated using the Lenti-X-Concentrator reagent (Takara Bio, catalog no. 631232) following the manufacturer’s instructions and resuspended in OPTI-MEM at 10% of the original culture volume. Lentiviral particles were subsequently aliquoted and frozen at −80°C.

### Lentivirus titration

K562 cells were seeded onto 24-well plate at 50k cells per well with 250ul fresh medium, supplemented with 16ug/ml polybrene. 250ul fresh medium was added to well 1 as the untransduced control. Wells 2-6 were transduced with lentivirus particles ranging from 250μl (1:1) to 15.625μl (1:16). Wells 7-12 were replicated with the same lentivirus mixture. The final concentration of polybrene was 8ul/ml. After culturing for 24 hours at 37°C, exchange the medium for 500ul fresh medium without polybrene. 72 hours post-transduction, flow cytometry (BD LSRFortessa) was employed to determine the ratio of transduced cells (TagBFP: sgRNA or mCherry: dCas9).

### Single cell sequencing

Wild type (WT) K562 cells were first transduced with desired amount (estimated based on the titration result) of the dCas9-KRAB lentivirus with polybrene (8 ug/ml) to achieve a high MOI (∼ 0.7, day -2). Two days later (day 0), they were infected with pooled sgRNA lentivirus (∼ 0.2 MOI) and polybrene. On day 3 post-transduction, cells were sorted using FACS for the mCherry (dCas9-KARB) and TagBFP (sgRNA) double-positive population. Sorted cells were cultured with fresh medium maintaining at a density of 100k cells/ml. Seven days after pooled sgRNA transduction (day 7), cells were sorted again for viability and double positive fluorescence, then harvested for sequencing immediately. Single cells were collected for the construction of 5’ single-cell gene expression (GEX) libraries and CRISPR guide RNA (gRNA) capture libraries using the Chromium X platform (10x Genomics, PN-1000287, PN-1000451, PN-1000263, PN-1000215). The libraries were prepared with a target recovery of approximately 10,000 cells. Both GEX and CRISPR libraries were sequenced on an Illumina NovaSeq 6000 system using a 100-cycle kit. The sequencing configuration included read 1 (28 bp), read 2 (10 bp), read 3 (10 bp), and read 4 (90 bp).

### Individual sgRNA cloning experiment

Eleven sgRNAs targeting 3 genes with 3 sgRNAs each and 2 non-targeting controls were individually cloned into pBA950 vector. After heat shock transformation, plasmids were extracted using the Miniprep kit (QIAGEN, catalog no. 27104). All plasmids were verified by XmnI (NEB, catalog no. R0194S) digest and Sanger sequencing with the primer 5’-CCTCGGCCTCTGCATAAATA-3’. dCas9-KRAB K562 cells were transfected with the eleven sgRNA plasmids in parallel. In each reaction, 100ul 1X Nucleofection buffer was prepared with 4.5ug plasmids. Following count, spin down 1.5 million cells and resuspend the pellet in the Nucleofection buffer with plasmids. Cells were then transferred to an electroporation cuvette and shocked using program T-016 (Amaxa Nucleofector I). When shock is over, quickly add 1ml media to the cuvette and plate the cells. 48 hours post-transfection, sort the TagBFP and mCherry double-positive cells population using FACS. 72 hours post-transfection, cells were harvested and stored at -80°C, awaiting subsequent RNA extraction.

### RT-qPCR

Total RNA was isolated using ReliaPrep™ RNA Cell Miniprep System (Promega, catalog no. Z6012) and converted to cDNA using SuperScipt IV Reverse Transcriptase kit (Invitrogen, catalog no. 18090200) following manufactures’ instructions. RT-qPCR reactions were prepared following Platinum™ Taq DNA Polymerase kit (Invitrogen, catalog no.10966034) supplemented with 1ul 1/2000 SYBR Green (Sigma, catalog no. S9430) in 25ul reactions. Target genes expression were determined with the primer pairs listed in Supplementary Table 4 and measured on a C1000 Touch Thermal Cycler (Bio-Rad). Relative expression of mRNA for different samples was determined by the ΔΔCT method using GAPDH as the internal reference, and visualized by Prism v8.0.2.

### Processing of single-cell sequencing data

Gene expression and sgRNA libraries were processed using the 10X Cell Ranger version 7.0.1 following default settings. Gene expression reads were aligned to the reference genome “refdata-gex-GRCh38-2020-A”. sgRNA reads were aligned to the reverse complemented sgRNA library using the pattern TTCCAGCTTAGCTCTTAAAC(BC). The “protospacer_calls_per_cell.csv” output file, generated through the application of the Gaussian Mixture Model within Cell Ranger, was used to assign sgRNAs to cells. For cells not listed in this file, they were either no guide molecules founded or lack of a confident call (didn’t pass the threshold). Cells with more than 1 feature (sgRNA) were filtered out.

Gene expression matrix were imported and analyzed using the Seurat package (version 4) in R. Cells were filtered for mitochondrial RNA percentage < 10%, unique genes detected > 400 and < 7000, and UMI counts < 60000. Finally, 6898 high-quality cells harboring single perturbations were used for all the further analysis. On-target knockdown (Fold change and P_val_adj) was calculated using the FindMarkers function in Seurat comparing the perturbed cells (cells containing sgRNAs against each target) to non-targeting cells (all the cells harboring non-targeting sgRNAs) with the parameter “pseudocount.use = 1e-9”. The identification of DEGs for each sgRNA was accomplished using FindMarkers with default settings comparing to cells harboring non-targeting sgRNAs.

### Mixscape analysis

We analyzed the aforementioned processed count matrix following the Mixscape pipeline (https://satijalab.org/seurat/articles/mixscape_vignette). To investigate the effect at the sgRNA level, we ran the RunMixscape (Mixscape command) with find.mode = TRUE and prtb.type = “KD”. The results of this Mixscape analysis were visualized using bar plots and further explored through a heatmap of differentially expressed genes (DEGs) across KD, NP, and NT cell subsets. These DEGs were identified by comparing KD cells to NT cells. To visualize perturbation-specific clusters, Linear Discriminant Analysis (LDA) was used as the dimensionality reduction method on NT cells and KD cells derived from efficient sgRNAs.

### Gene regulatory networks inference and validation

Gene regulatory networks inference was started by first running GENIE3^68^ (SCENIC command) to identify co-expression modules. To enhance the accuracy of our results, we included all 6,898 cells with at least one perturbation, as incorporating multiple cell conditions improves module detection. In this step, only positive correlation were kept and used for the further analysis. Next, we used RcisTarget^48^ (SCENIC command) to identify regulons, consisting of transcription factors (TFs) and their direct-binding targets. This analysis initially identified 307 regulons. However, 189 of these were labeled as “extended,” indicating low-confidence motif detection. To ensure robustness, we excluded these low-confidence regulons, retaining 118 high-confidence regulons for further analysis.

Next, we calculated regulon activity in each cell using AUCell^48^ (SCENIC command) and generated heatmaps at both the sgRNA level and gene level to visualize activity patterns. At the gene level, the MYB KD group included all KD cells from three sgRNAs, the LMO2 KD group consisted of KD cells from LMO2-1 and LMO2-3, and the GATA2 KD group contained KD cells from GATA2-3. All other cells were categorized as NP and labeled as MYB NP, LMO2 NP, and GATA2 NP, respectively. Then we calculated the regulon-specific score (RSS) to filter out regulons that exhibited high activity across all clusters, as these were not of interest. Instead, we focused on regulons that displayed high activity in specific clusters, allowing us to identify key regulatory differences among the KD groups. To incorporate low-activity regulons, we applied a filter of RAC < -1.

After combining low-activity regulons (RAC < -1) and high-specificity regulons (RSS Z-score > 0.5) for each cluster (MYB KD, GATA2 KD, and LMO2 KD), we observed that some regulons were shared between two clusters, suggesting potential regulatory links between the corresponding KD genes. However, one limitation of this approach is its reliance on regulon activity measurements, which rank all genes in a regulon within each cell. This can lead to false positive regulons, where high regulon activity is driven by a few highly expressed genes that do not include the transcription factor (TF) itself. Since all genes in a regulon should be direct TF targets, the TF’s expression level is expected to change as well. To address this issue, we calculated Fold Change (FC) for the TF in each selected regulon and only considered regulons with TF |log2 FC| > 0.5 as potentially relevant regulons.

To validate the GRNs, we selected some genes from regulons with |log2 FC| > 0.5 and adjusted p-value < 0.05. We then performed knockdown (KD) experiments like “Individual sgRNA cloning experiment”. To ensure robust comparisons, we pooled cDNA from LMO2-1 and LMO2-3 as the LMO2 KD group, and GATA2-3 as the GATA2 KD group. RT-qPCR analysis was conducted to assess the expression changes of the selected validation genes.

## Published dataset analysis

Raw data from two published papers, Wu et al. (2022) and Joseph M.R et al. (2022), were retrieved from the Gene Expression Omnibus (GEO) under the accession codes GSE150062 and GSM6210117, respectively. They were analyzed following the same approach we applied for the aforementioned TFs described with headers: “Processing of single-cell sequencing data” and “Mixscape analysis”.

## Author contribution

HZ and EM conceived the study. HZ performed the experiments and analyzed the data under the supervision of EM. EB performed the single-cell experiments. PZ assisted with experimental work and data analysis. HZ and EM wrote the manuscript. All authors read and approved the final version.

## Data availability

Raw and processed data have been deposited in NCBI-GEO under accession number GSE308682, and are available upon publication.

## Code availability

The code used for the data analysis is based on publicly available pipelines. The Mixscape pipeline can be accessed at https://satijalab.org/seurat/articles/mixscape_vignette, and the SCENIC pipeline is available on GitHub at https://github.com/aertslab/SCENIC.

## Acknowledgements

We thank the China Scholarship Council for financially supporting P.Z., and H.Z. We would like to acknowledge Lucas Kuijpers for helpful discussions. We thank Tsung Wai Kan for assistance with FACS sorting of fluorescent cells. Wilfred van Ijcken and Zakia Azmani are gratefully acknowledged for designing and performing the next-generation sequencing used for library quality control.

## Ethics declaration

All authors declare that they have no conflicts of interest.

## Supplementary Figures legends

**Supplementary Figure 1. Quality control of the pooled sgRNA library.**

(A) Number of reads detected for each sgRNA in the plasmid pool. (B) Number of reads detected after amplification from genomic DNA of cells transduced with the pooled lentiviral library. To indicate distribution of sgRNAs in the library, Fold Change (number of counts for the 90th percentile of sgRNAs detected / number of counts for the 10th percentile of sgRNAs detected) in each quality control step was calculated.

**Supplementary Figure 2. Expression level of genes targeted for inactivation**

Target gene expression level for different cell groups at gene level (A-F) and sgRNA level (G-L). NT: Cells with non-targeting sgRNAs. LMO2-KD: Cells with all the three LMO2 sgRNAs. LMO2-1: Cells with LMO2 sgRNA-1. LMO2-2: Cells with LMO2 sgRNA-2. LMO2-3: Cells with LMO2 sgRNA-3. LDB1-KD: Cells with all the three LDB1 sgRNAs. LMO2-1: Cells with LDB1 sgRNA-1. LDB1-2: Cells with LDB1 sgRNA-2. LDB1-3: Cells with LDB1 sgRNA-3. GATA2-KD: Cells with all the three GATA2 sgRNAs. LMO2-1: Cells with GATA2 sgRNA-1. GATA2-2: Cells with GATA2 sgRNA-2. GATA2-3: Cells with GATA2 sgRNA-3.

RNUX1-KD: Cells with all the three RNUX1 sgRNAs. LMO2-1: Cells with RNUX1 sgRNA-1. RNUX1-2: Cells with RNUX1 sgRNA-2. RNUX1-3: Cells with RNUX1 sgRNA-3. TCF3-KD: Cells with all the three TCF3 sgRNAs. LMO2-1: Cells with TCF3 sgRNA-1. TCF3-2: Cells with TCF3 sgRNA-2. TCF3-3: Cells with TCF3 sgRNA-3. FLI1-KD: Cells with all the three FLI1 sgRNAs. LMO2-1: Cells with FLI1 sgRNA-1. FLI1-2: Cells with FLI1 sgRNA-2. FLI1-3: Cells with FLI1 sgRNA-3.

**Supplementary Figure 3. Heatmap of DEGs after LMO2 & GATA2 Knock Down**

(A) Heatmap of DEGs identified by comparing LMO2 knock-down and non-perturbed cells (as estimated by Mixscape) to non-targeting cells. (B) Heatmap of DEGs identified by comparing GATA2 knock-down and non-perturbed cells (as estimated by Mixscape) to non-targeting cells.

**Supplementary Figure 4. Validation of differentially expressed genes after gene KD**

Relative expression of GATA2 (A) and LMO2 (B) after gene perturbation using CRISPRi. Each sgRNA was transfected into dCas9 K562 cells separately, then RT-qPCR was performed independently. (C) Relative expression of 5 up regulated genes after transduction of single sgRNAs targeting GATA2. (D) Relative expression of 5 down regulated genes after transduction of single sgRNAs targeting GATA2. (E) Relative expression of 5 up regulated genes after transduction of single sgRNAs targeting LMO2. (F) Relative expression of 5 down regulated genes after transduction of single sgRNAs targeting LMO2. Data are presented as mean ± SD. Statistical significance was determined using two-tailed Student’s t-test.

Significance levels are indicated as follows: ns, not significant; * p < 0.05; ** p < 0.01; *** p < 0.001; **** p < 0.0001.

**Supplementary Figure 5. Gene enrichment analysis and MYB occupancy status at TSS of DEGs**

(A) Gene set enrichment analysis of DEGs following MYB knockdown (KD) for up and down regulated genes. (B) MYB ChIP-seq peak occupancy in K562 cells centered around transcription start sites (TSS, position 0) of DEGs. Up: all the up-regulated DEGs we identified after MYB KD. Down: all the down-regulated DEGs we identified after MYB KD. Control: K562 cells expressing empty vector. Each condition represents the average signal across three biological replicates.

**Supplementary Figure 6. Gene enrichment analysis after GATA2 KD and GATA2 occupancy at TSS of DEGs**

(A) Gene set enrichment analysis of DEGs following GATA2 KD . (B) GATA2 ChIP-seq peak occupancy in K562 cells centered around transcription start sites (TSS, position 0) of DEGs. FoldChange of ChIP-seq signal comparing GATA2 to control sample. Up: all the up-regulated DEGs we identified after GATA2 KD. Down: all the down-regulated DEGs we identified after GATA2 KD. Control: K562 cells expressing empty vector.

**Supplementary Figure 7. Regulon activity at the sgRNA level and RT-qPCR validation of inferred GRNs.**

(A) Regulon activity across individual sgRNA level. The X-axis represents cell identities (including NT, a combination of five non-targeting sgRNAs), and the Y-axis shows regulons. (B, C) RT-qPCR validation of representative regulon target genes following knockdown of their predicted regulators. Data are presented as mean ± SD. Statistical significance was determined using two-tailed Student’s t-test. Significance levels are indicated as follows: ns, not significant; * p < 0.05; ** p < 0.01; *** p < 0.001; **** p < 0.0001.

**Supplementary Figure 8. Analysis of two published Perturb-seq CRISPRi datasets.**

(A, B) Percentage of knockdown (KD) sgRNAs identified in each dataset. (C, D) UMAP visualizations of cells targeted by KD-like sgRNAs in the two datasets. (E, F) Overlap of sgRNAs passing both fold-change and Mixscape thresholds.

**Supplementary Figure 9. FACS for mCherry-positive K562 cells.**

mCherry fluorescence were measured for WT K562 cells (A) and dCas9 transduced K562 cells (B). WT: WT K562 cells. D9: dCas9 transduced K562 cells. Only mCherry-positive cells were collected.

## Supplementary Tables descriptions

**Supplementary Table 1.** Differentially expressed genes identified using KD cells comparing to NT cells.

**Supplementary Table 2.** Important regulons screened using the regulon activity score (RAC) and regulon specificity score (RSS).

**Supplementary Table 3.** sgRNA protospacers used in this study and 72/73 bp ssDNA oligos ordered from IDT.

**Supplementary Table 4.** RT-qPCR primers used in this study.

## Reference

1 Schraivogel, D., Steinmetz, L. M. & Parts, L. Pooled Genome-Scale CRISPR Screens in Single Cells. Annu Rev Genet 57, 223–244 (2023). 10.1146/annurev-genet-072920-013842

2 Dixit, A. et al. Perturb-Seq: Dissecting Molecular Circuits with Scalable Single-Cell RNA Profiling of Pooled Genetic Screens. Cell 167, 1853–1866.e1817 (2016). 10.1016/j.cell.2016.11.038

3 Datlinger, P. et al. Pooled CRISPR screening with single-cell transcriptome readout. Nat Methods 14, 297–301 (2017). 10.1038/nmeth.4177

4 Agrotis, A. & Ketteler, R. A new age in functional genomics using CRISPR/Cas9 in arrayed library screening. Front Genet 6, 300 (2015). 10.3389/fgene.2015.00300

5 Shalem, O. et al. Genome-scale CRISPR-Cas9 knockout screening in human cells. Science 343, 84–87 (2014). 10.1126/science.1247005

6 Adamson, B. et al. A Multiplexed Single-Cell CRISPR Screening Platform Enables Systematic Dissection of the Unfolded Protein Response. Cell 167, 1867–1882.e1821 (2016). 10.1016/j.cell.2016.11.048

7 Jaitin, D. A. et al. Dissecting Immune Circuits by Linking CRISPR-Pooled Screens with Single-Cell RNA-Seq. Cell 167, 1883–1896.e1815 (2016). 10.1016/j.cell.2016.11.039

8 So, R. W. L. et al. Application of CRISPR genetic screens to investigate neurological diseases. Mol Neurodegener 14, 41 (2019). 10.1186/s13024-019-0343-3

9 Wang, T., Wei, J. J., Sabatini, D. M. & Lander, E. S. Genetic screens in human cells using the CRISPR-Cas9 system. Science 343, 80–84 (2014). 10.1126/science.1246981

10 Shifrut, E. et al. Genome-wide CRISPR Screens in Primary Human T Cells Reveal Key Regulators of Immune Function. Cell 175, 1958–1971.e1915 (2018). 10.1016/j.cell.2018.10.024

11 Henriksson, J. et al. Genome-wide CRISPR Screens in T Helper Cells Reveal Pervasive Crosstalk between Activation and Differentiation. Cell 176, 882–896.e818 (2019). 10.1016/j.cell.2018.11.044

12 Guerriero, M. L. et al. Delivering Robust Candidates to the Drug Pipeline through Computational Analysis of Arrayed CRISPR Screens. SLAS Discov 25, 646–654 (2020). 10.1177/2472555220921132

13 Kim, H. S. et al. Arrayed CRISPR screen with image-based assay reliably uncovers host genes required for coxsackievirus infection. Genome Res 28, 859–868 (2018). 10.1101/gr.230250.117

14 Ross-Thriepland, D. et al. Arrayed CRISPR Screening Identifies Novel Targets That Enhance the Productive Delivery of mRNA by MC3-Based Lipid Nanoparticles. SLAS Discov 25, 605–617 (2020). 10.1177/2472555220925770

15 Blomen, V. A. et al. Gene essentiality and synthetic lethality in haploid human cells. Science 350, 1092–1096 (2015). 10.1126/science.aac7557

16 Marceau, C. D. et al. Genetic dissection of Flaviviridae host factors through genome-scale CRISPR screens. Nature 535, 159–163 (2016). 10.1038/nature18631

17 Porichis, F. et al. High-throughput detection of miRNAs and gene-specific mRNA at the single-cell level by flow cytometry. Nat Commun 5, 5641 (2014). 10.1038/ncomms6641

18 Haney, M. S. et al. Identification of phagocytosis regulators using magnetic genome-wide CRISPR screens. Nat Genet 50, 1716–1727 (2018). 10.1038/s41588-018-0254-1

19 Fulco, C. P. et al. Activity-by-contact model of enhancer-promoter regulation from thousands of CRISPR perturbations. Nat Genet 51, 1664–1669 (2019). 10.1038/s41588-019-0538-0

20 Song, Q. et al. Direct-seq: programmed gRNA scaffold for streamlined scRNA-seq in CRISPR screen. Genome Biol 21, 136 (2020). 10.1186/s13059-020-02044-w

21 McFarland, J. M. et al. Multiplexed single-cell transcriptional response profiling to define cancer vulnerabilities and therapeutic mechanism of action. Nat Commun 11, 4296 (2020). 10.1038/s41467-020-17440-w

22 Yan, X. et al. High-content imaging-based pooled CRISPR screens in mammalian cells. J Cell Biol 220 (2021). 10.1083/jcb.202008158

23 Schraivogel, D. et al. High-speed fluorescence image-enabled cell sorting. Science 375, 315–320 (2022). 10.1126/science.abj3013

24 Li, Y. et al. UDA-seq: universal droplet microfluidics-based combinatorial indexing for massive-scale multimodal single-cell sequencing. Nat Methods 22, 1199–1212 (2025). 10.1038/s41592-024-02586-y

25 Metzner, E., Southard, K. M. & Norman, T. M. Multiome Perturb-seq unlocks scalable discovery of integrated perturbation effects on the transcriptome and epigenome. Cell Syst 16, 101161 (2025). 10.1016/j.cels.2024.12.002

26 Papalexi, E. et al. Characterizing the molecular regulation of inhibitory immune checkpoints with multimodal single-cell screens. Nat Genet 53, 322–331 (2021). 10.1038/s41588-021-00778-2

27 Yan, R. E. et al. Pooled CRISPR screens with joint single-nucleus chromatin accessibility and transcriptome profiling. Nat Biotechnol (2024). 10.1038/s41587-024-02475-x

28 Replogle, J. M. et al. Combinatorial single-cell CRISPR screens by direct guide RNA capture and targeted sequencing. Nat Biotechnol 38, 954–961 (2020). 10.1038/s41587-020-0470-y

29 Schmidt, R. et al. CRISPR activation and interference screens decode stimulation responses in primary human T cells. Science 375, eabj4008 (2022). 10.1126/science.abj4008

30 Legut, M. et al. A genome-scale screen for synthetic drivers of T cell proliferation. Nature 603, 728–735 (2022). 10.1038/s41586-022-04494-7

31 Horlbeck, M. A. et al. Compact and highly active next-generation libraries for CRISPR-mediated gene repression and activation. Elife 5 (2016). 10.7554/eLife.19760

32 Boettcher, M. & McManus, M. T. Choosing the Right Tool for the Job: RNAi, TALEN, or CRISPR. Mol Cell 58, 575–585 (2015). 10.1016/j.molcel.2015.04.028

33 Tian, R. et al. CRISPR Interference-Based Platform for Multimodal Genetic Screens in Human iPSC-Derived Neurons. Neuron 104, 239–255.e212 (2019). 10.1016/j.neuron.2019.07.014

34 Wu, D. et al. Dual genome-wide coding and lncRNA screens in neural induction of induced pluripotent stem cells. Cell Genom 2 (2022). 10.1016/j.xgen.2022.100177

35 Truchi, M. et al. Detecting subtle transcriptomic perturbations induced by lncRNAs knock-down in single-cell CRISPRi screening using a new sparse supervised autoencoder neural network. Front Bioinform 4, 1340339 (2024). 10.3389/fbinf.2024.1340339

36 Chen, B. et al. Dynamic imaging of genomic loci in living human cells by an optimized CRISPR/Cas system. Cell 155, 1479–1491 (2013). 10.1016/j.cell.2013.12.001

37 Tyner, J. W. et al. Functional genomic landscape of acute myeloid leukaemia. Nature 562, 526–531 (2018). 10.1038/s41586-018-0623-z

38. Comprehensive genomic characterization defines human glioblastoma genes and core pathways. Nature 455, 1061–1068 (2008). 10.1038/nature07385

39 Sanson, K. R. et al. Optimized libraries for CRISPR-Cas9 genetic screens with multiple modalities. Nat Commun 9, 5416 (2018). 10.1038/s41467-018-07901-8

40 Lemma, R. B. et al. Chromatin occupancy and target genes of the haematopoietic master transcription factor MYB. Sci Rep 11, 9008 (2021). 10.1038/s41598-021-88516-w

41 Fuglerud, B. M. et al. A c-Myb mutant causes deregulated differentiation due to impaired histone binding and abrogated pioneer factor function. Nucleic Acids Res 45, 7681–7696 (2017). 10.1093/nar/gkx364

42 Wang, X., Angelis, N. & Thein, S. L. MYB - A regulatory factor in hematopoiesis. Gene 665, 6–17 (2018). 10.1016/j.gene.2018.04.065

43 Peters, I. J. A., de Pater, E. & Zhang, W. The role of GATA2 in adult hematopoiesis and cell fate determination. Front Cell Dev Biol 11, 1250827 (2023). 10.3389/fcell.2023.1250827

44 Zhu, H. et al. Regulation of the lmo2 promoter during hematopoietic and vascular development in zebrafish. Dev Biol 281, 256–269 (2005). 10.1016/j.ydbio.2005.01.034

45 Zuber, J. et al. RNAi screen identifies Brd4 as a therapeutic target in acute myeloid leukaemia. Nature 478, 524–528 (2011). 10.1038/nature10334

46 Lorenzo, P. I. et al. Identification of c-Myb Target Genes in K562 Cells Reveals a Role for c-Myb as a Master Regulator. Genes Cancer 2, 805–817 (2011). 10.1177/1947601911428224

47. An integrated encyclopedia of DNA elements in the human genome. Nature 489, 57–74 (2012). 10.1038/nature11247

48 Aibar, S. et al. SCENIC: single-cell regulatory network inference and clustering. Nature methods 14, 1083–1086 (2017).

49 Fagnan, A., Drissen, R., Di Genua, C., Meng, Y. & Nerlov, C. Myelo-Erythroid Lineage Segregation Is Regulated By the GATA-2 Interactome. Blood 142, 2687 (2023). 10.1182/blood-2023-190285

50 Schmidt, M., Nazarov, V., Stevens, L., Watson, R. & Wolff, L. Regulation of the resident chromosomal copy of c-myc by c-Myb is involved in myeloid leukemogenesis. Mol Cell Biol 20, 1970–1981 (2000). 10.1128/mcb.20.6.1970-1981.2000

51 Lohmann, F. & Bieker, J. J. Activation of Eklf expression during hematopoiesis by Gata2 and Smad5 prior to erythroid commitment. Development 135, 2071–2082 (2008). 10.1242/dev.018200

52 Lu, L. et al. LMO2 promotes the development of AML through interaction with transcription co-regulator LDB1. Cell Death Dis 14, 518 (2023). 10.1038/s41419-023-06039-w

53 Menendez-Gonzalez, J. B. et al. Gata2 as a Crucial Regulator of Stem Cells in Adult Hematopoiesis and Acute Myeloid Leukemia. Stem Cell Reports 13, 291–306 (2019). 10.1016/j.stemcr.2019.07.005

54 Inoue, A. et al. Elucidation of the role of LMO2 in human erythroid cells. Exp Hematol 41, 1062–1076.e1061 (2013). 10.1016/j.exphem.2013.09.003

55 Pan, Y., Tian, R., Lee, C., Bao, G. & Gibson, G. Fine-mapping within eQTL credible intervals by expression CROP-seq. Biol Methods Protoc 5, bpaa008 (2020). 10.1093/biomethods/bpaa008

56 Greatbatch, C. J. et al. High throughput functional profiling of genes at intraocular pressure loci reveals distinct networks for glaucoma. Hum Mol Genet 33, 739–751 (2024). 10.1093/hmg/ddae003

57 Zhao, X., Li, J., Liu, Z. & Powers, S. Combinatorial CRISPR/Cas9 Screening Reveals Epistatic Networks of Interacting Tumor Suppressor Genes and Therapeutic Targets in Human Breast Cancer. Cancer Res 81, 6090–6105 (2021). 10.1158/0008-5472.Can-21-2555

58 Replogle, J. M. et al. Maximizing CRISPRi efficacy and accessibility with dual-sgRNA libraries and optimal effectors. Elife 11 (2022). 10.7554/eLife.81856

59 Dean, A. PU.1 chromosomal dynamics are linked to LDB1. Blood 132, 2615–2616 (2018). 10.1182/blood-2018-10-880781

60 Goyama, S. et al. Transcription factor RUNX1 promotes survival of acute myeloid leukemia cells. J Clin Invest 123, 3876–3888 (2013). 10.1172/jci68557

61 Kotmayer, L. et al. GATA2 deficiency and MDS/AML: Experimental strategies for disease modelling and future therapeutic prospects. Br J Haematol 199, 482–495 (2022). 10.1111/bjh.18330

62 Lin, X. C. et al. Molecular dysfunctions in acute myeloid leukemia revealed by integrated analysis of microRNA and transcription factor. Int J Oncol 48, 2367–2380 (2016). 10.3892/ijo.2016.3489

63 Takao, S. et al. Convergent organization of aberrant MYB complex controls oncogenic gene expression in acute myeloid leukemia. Elife 10 (2021). 10.7554/eLife.65905

64 Lee, L. M. et al. A novel network pharmacology approach for leukaemia differentiation therapy using Mogrify(®). Oncogene 41, 5160–5175 (2022). 10.1038/s41388-022-02505-5

65 Ikonomi, P. et al. Levels of GATA-1/GATA-2 transcription factors modulate expression of embryonic and fetal hemoglobins. Gene 261, 277–287 (2000). 10.1016/s0378-1119(00)00510-2

66 Suzuki, M. et al. GATA factor switching from GATA 2 to GATA 1 contributes to erythroid differentiation. Genes to Cells 18, 921–933 (2013).

67 Nandakumar, S. K. et al. Low-level GATA2 overexpression promotes myeloid progenitor self-renewal and blocks lymphoid differentiation in mice. Experimental hematology 43, 565–577. e510 (2015).

68 Huynh-Thu, V. A., Irrthum, A., Wehenkel, L. & Geurts, P. Inferring regulatory networks from expression data using tree-based methods. PloS one 5, e12776 (2010).

