## Supplementary figures and images for "Insights from pooled CRISPRi single-cell screens in K562 cells reveal gene functions, regulatory networks, and highlight opportunities and limitations"

### Supplementary Figure 1

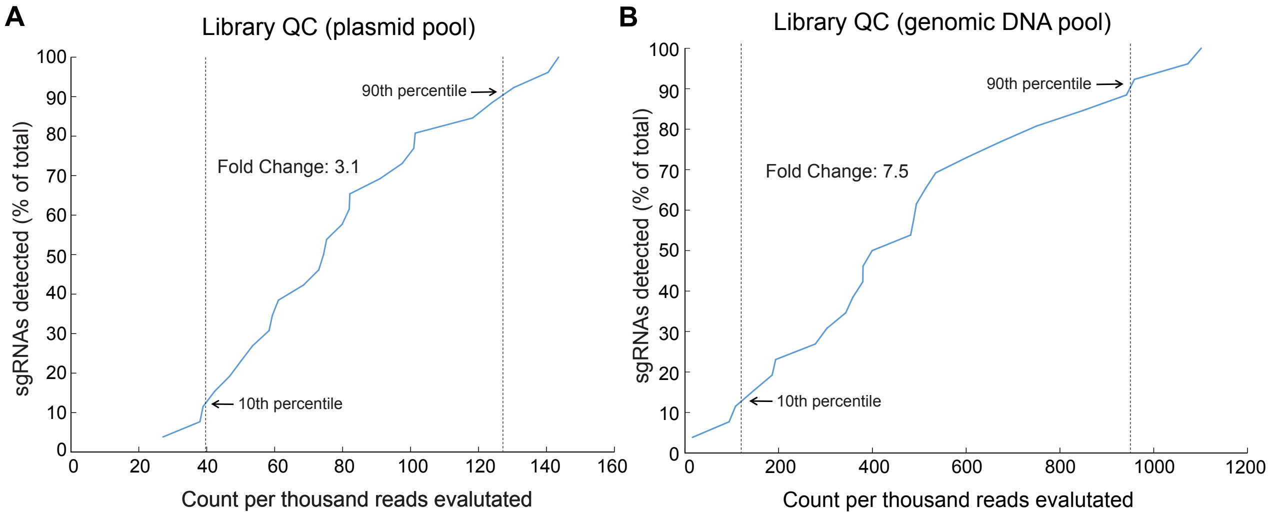


Zhang et al. - 2025

### Supplementary Figure 2

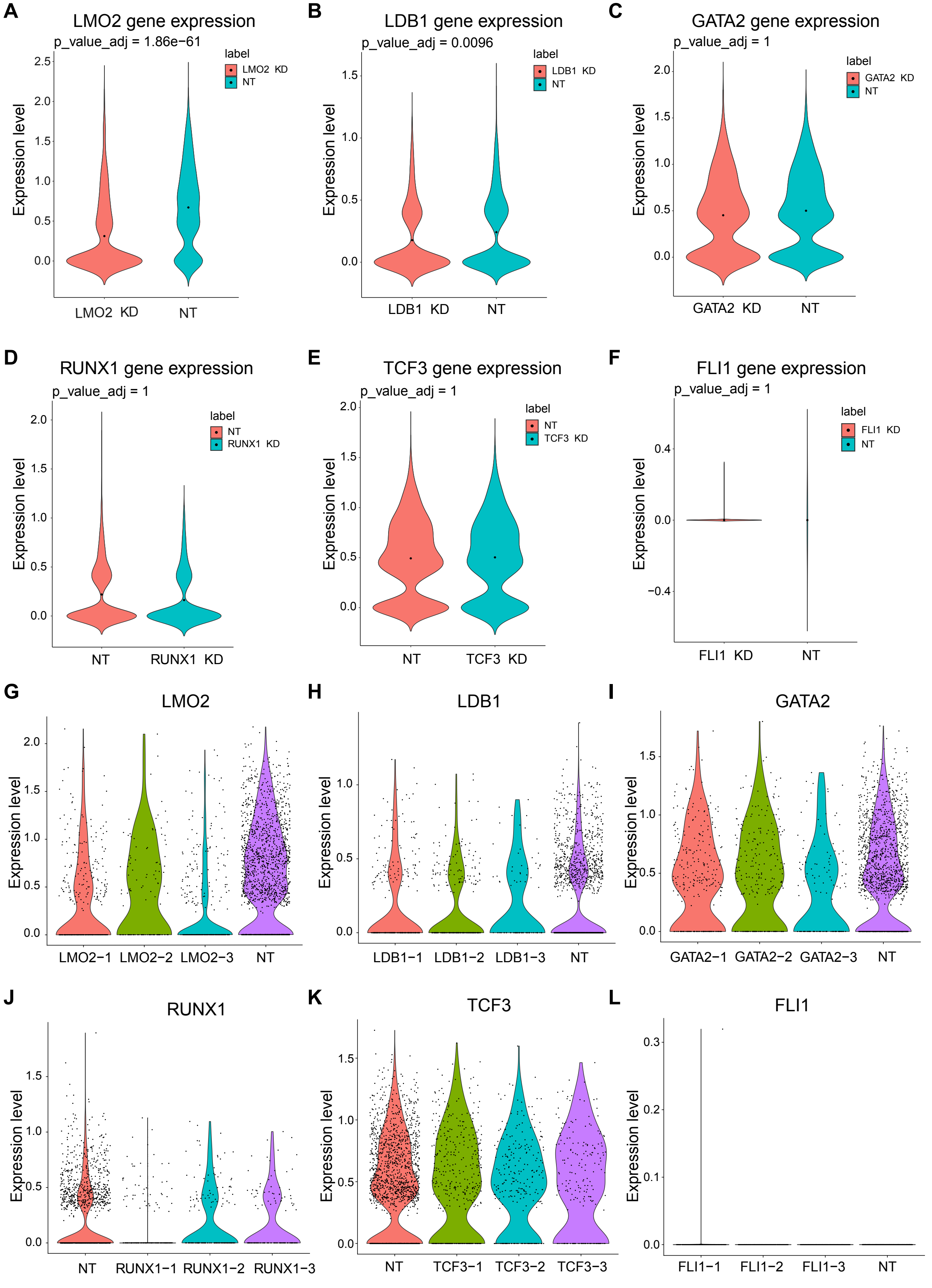


Zhang et al. - 2025

### Supplementary Figure 3

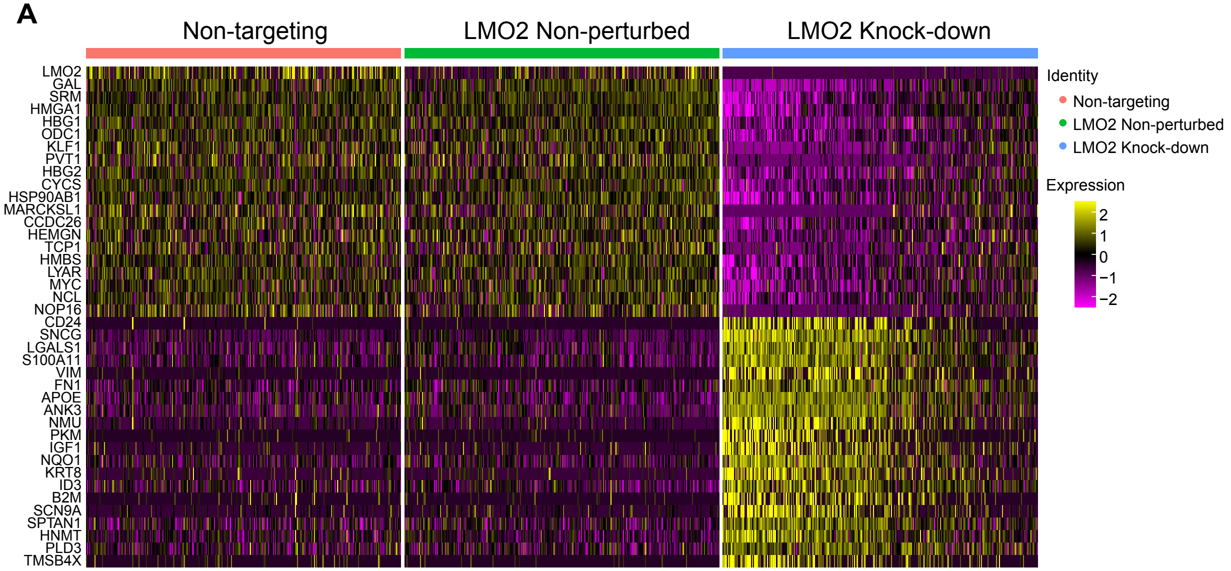

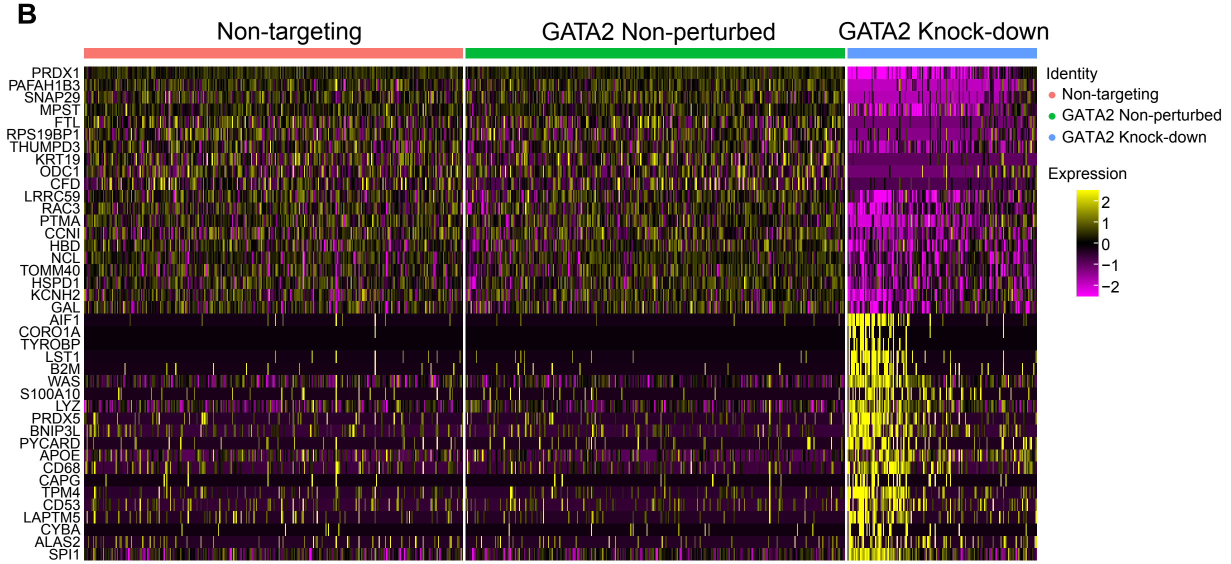

### Supplementary Figure 4

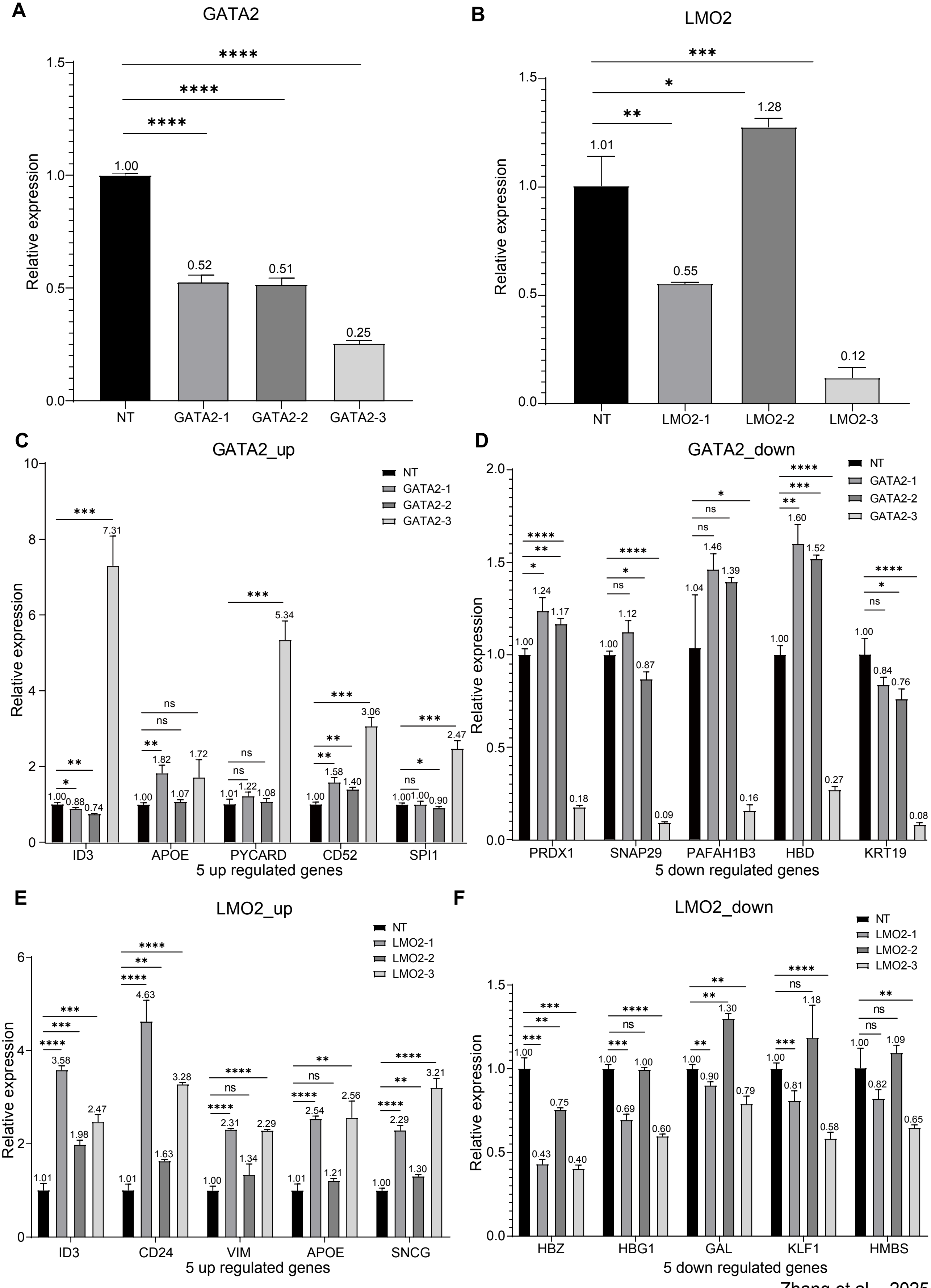


Zhang et al. - 2025

### Supplementary Figure 5

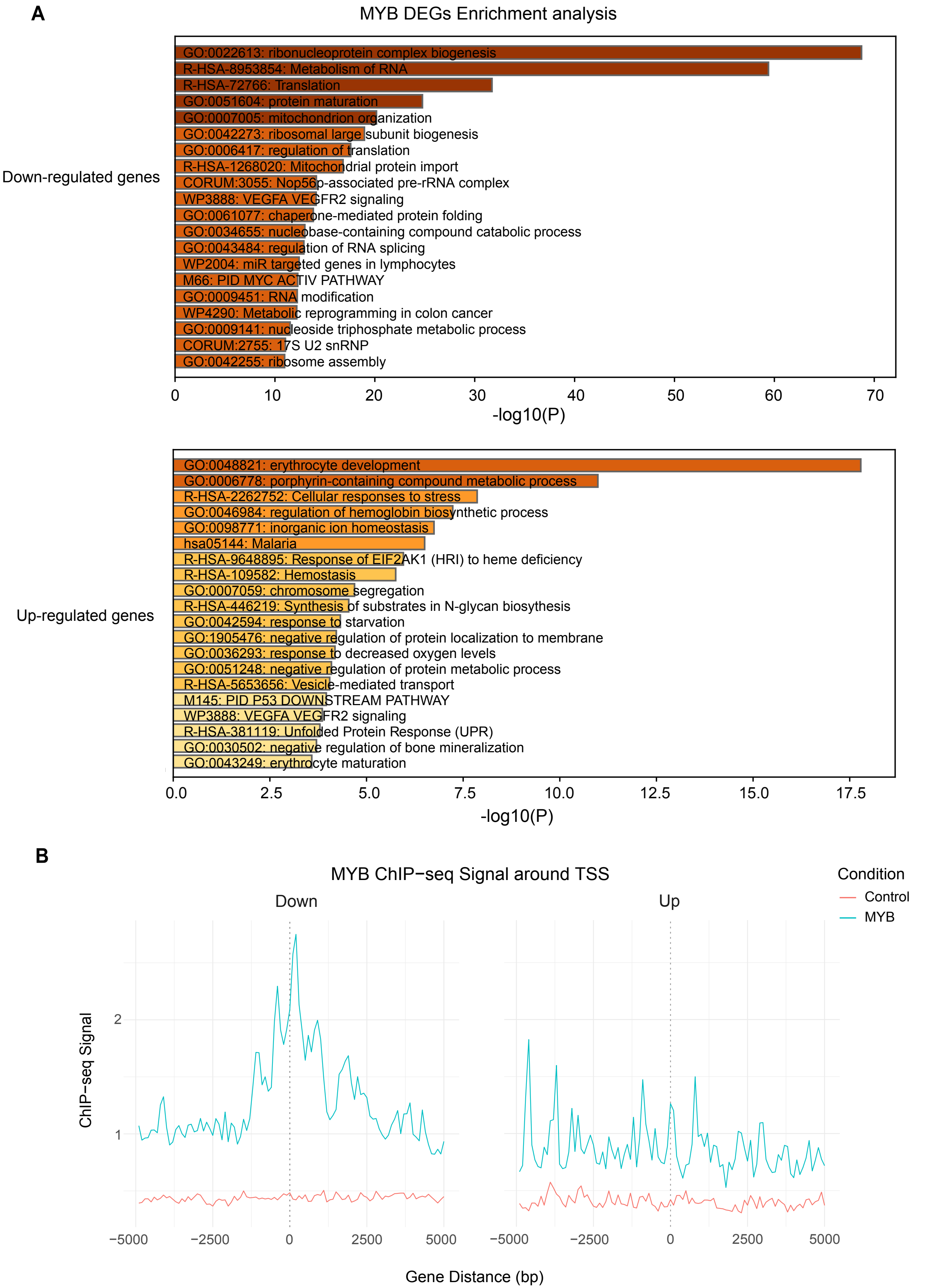


Zhang et al. - 2025

### Supplementary Figure 6

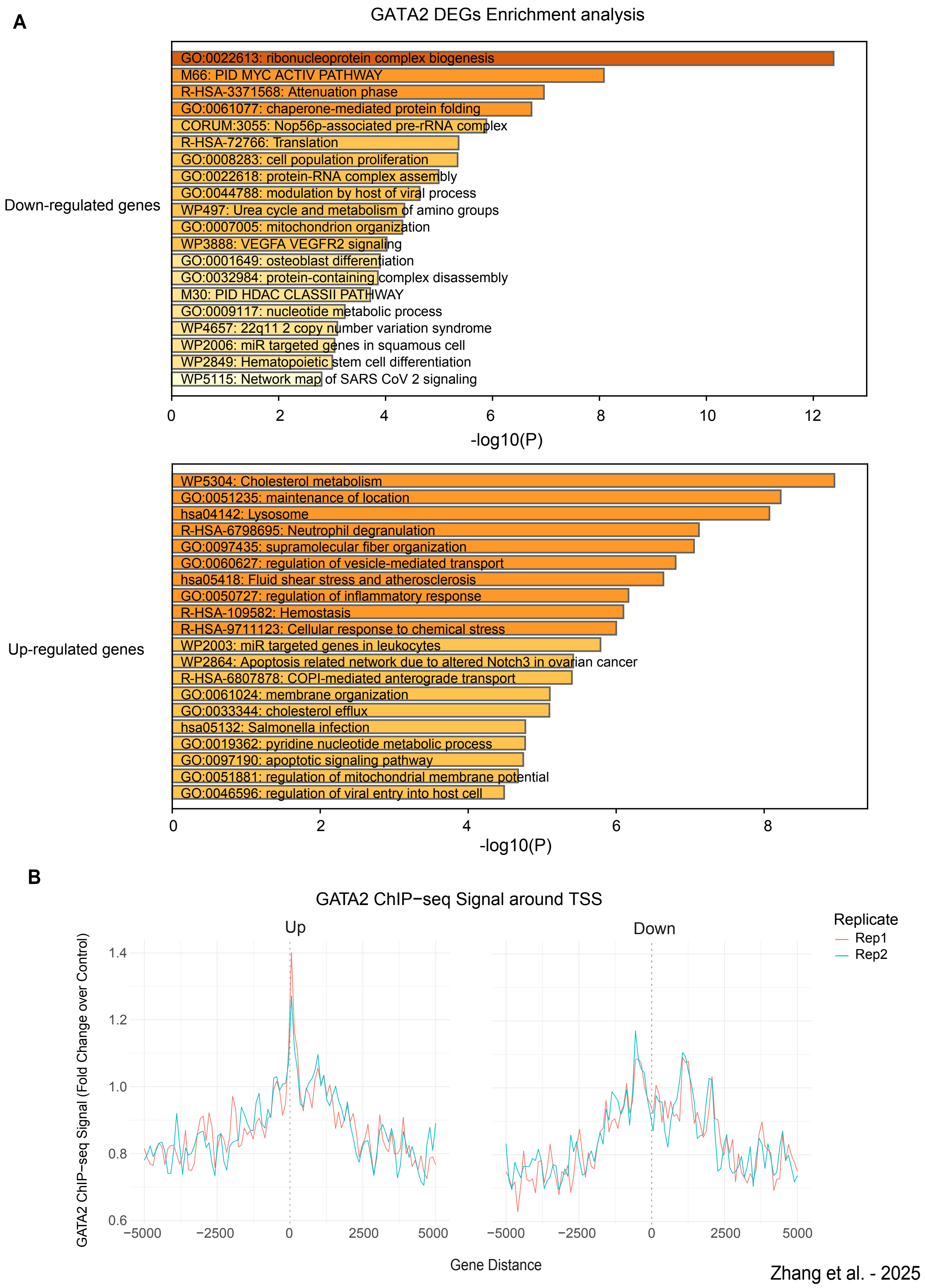

### Supplementary Figure 7

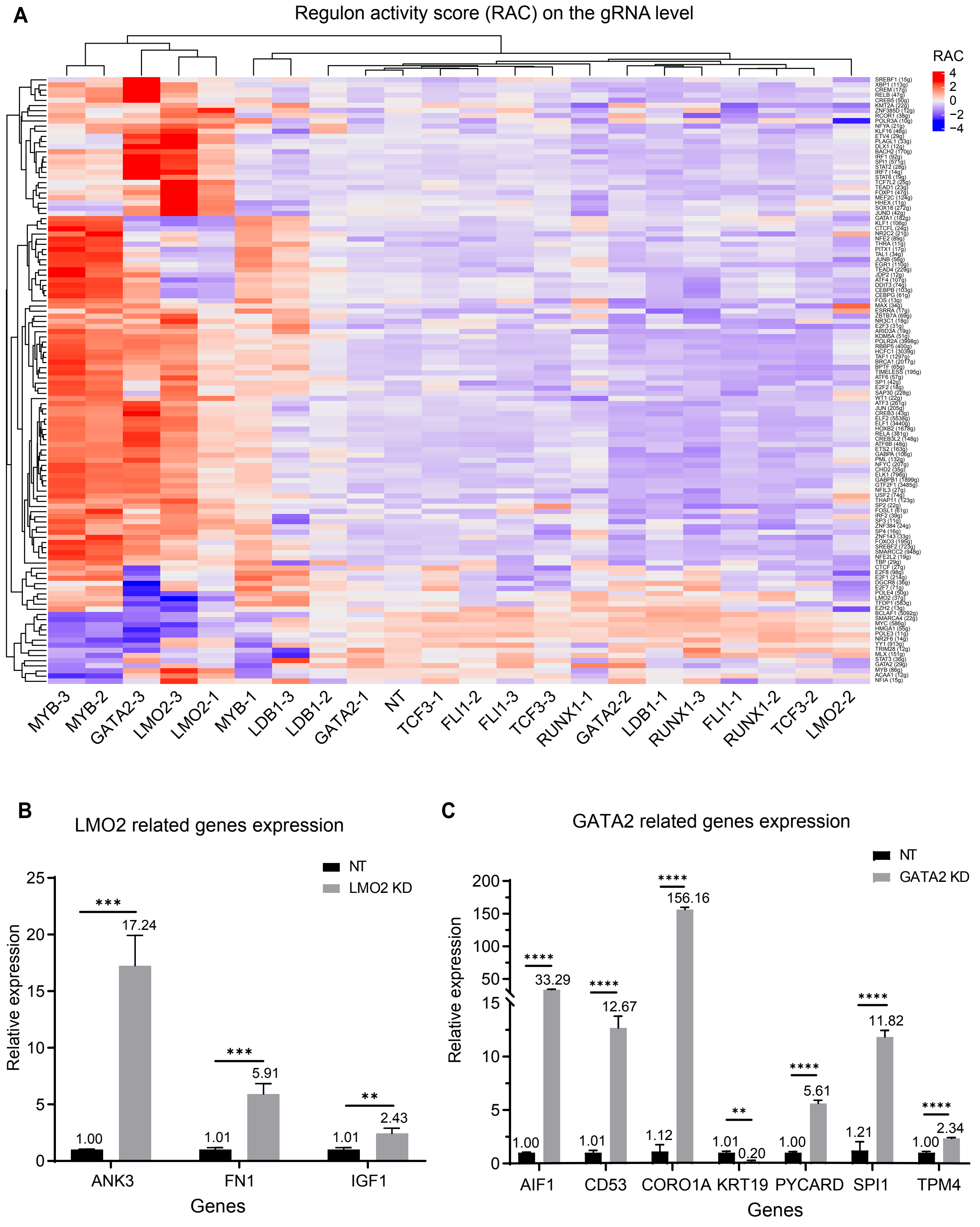


Zhang et al. - 2025

### Supplementary Figure 8

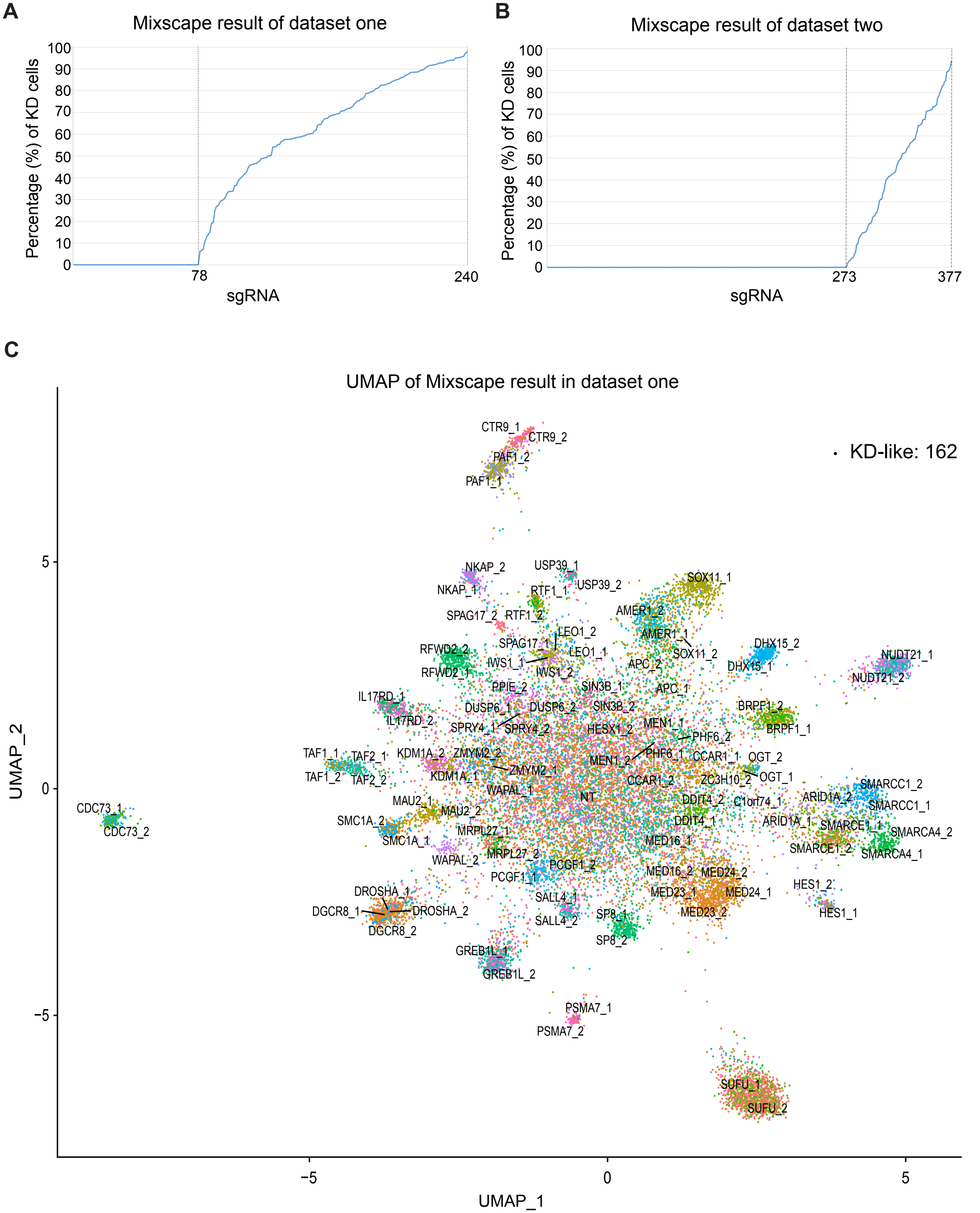


Zhang et al. - 2025

### Supplementary Figure 8_2

**D**

UMAP of Mixscape result in dataset two

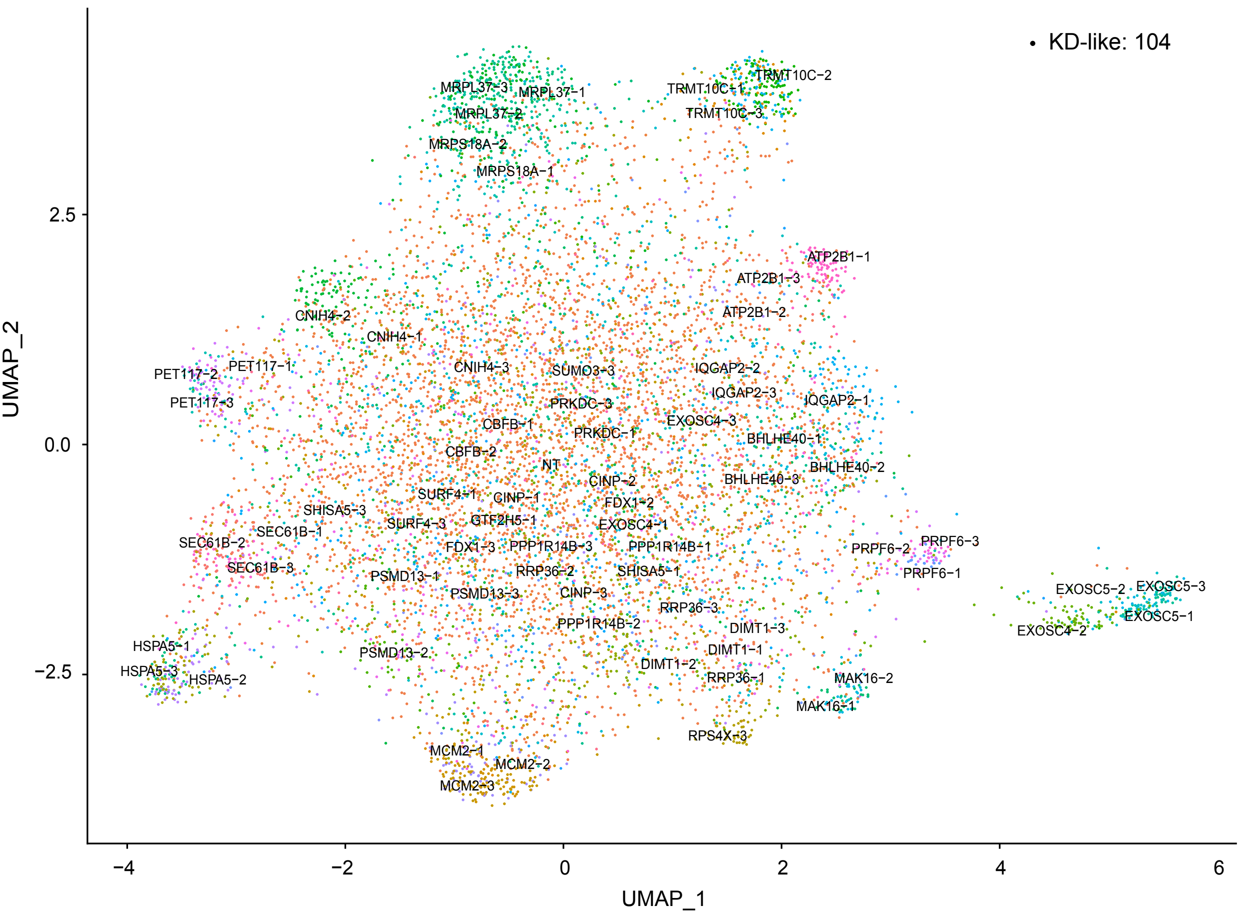


**E**

KD sgRNA in the dataset one

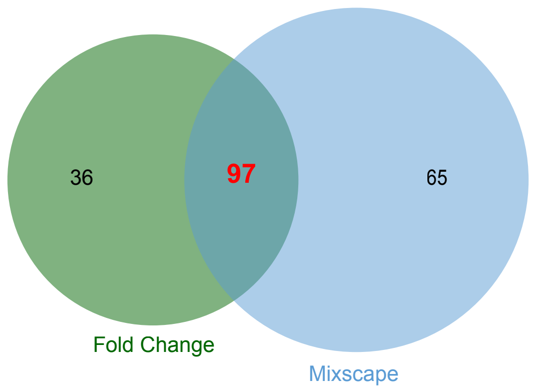


**F**

KD sgRNA in the dataset two

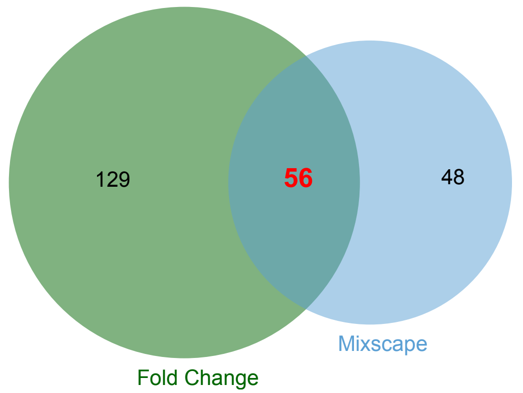
