## Supplementary Figure 9 for "Insights from pooled CRISPRi single-cell screens in K562 cells reveal gene functions, regulatory networks, and highlight opportunities and limitations"

**A** WT K562

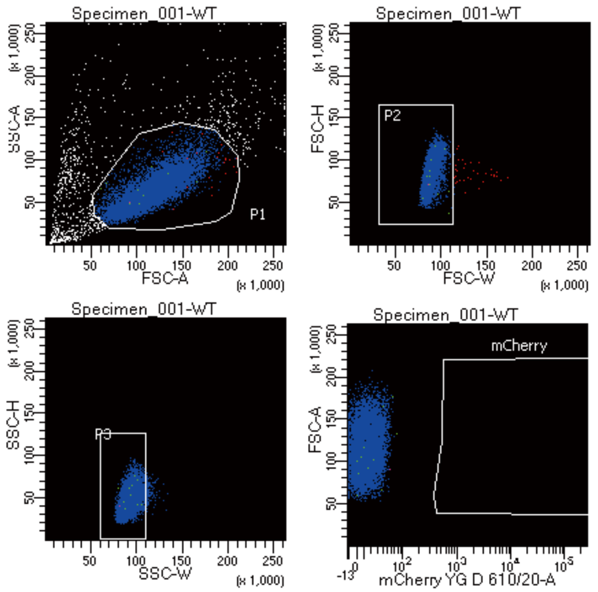


Tube: WT

| Population | #Events | %Parent | %Total |
|---|---|---|---|
| All Events | 10,000 | #### | 100.0 |
| P1 | 9,123 | 91.2 | 91.2 |
| P2 | 9,090 | 99.6 | 90.9 |
| P3 | 8,989 | 98.9 | 89.9 |
| mCherry | 0 | 0.0 | 0.0 |

**B** dCas9 transduced K562

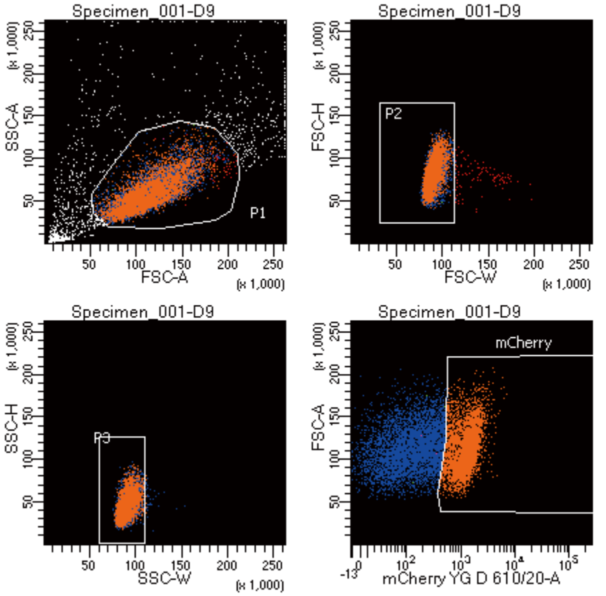


Tube: D9

| Population | #Events | %Parent | %Total |
|---|---|---|---|
| All Events | 10,000 | #### | 100.0 |
| P1 | 8,684 | 86.8 | 86.8 |
| P2 | 8,588 | 98.9 | 85.9 |
| P3 | 8,558 | 99.7 | 85.6 |
| mCherry | 4,172 | 48.7 | 41.7 |

Zhang et al. - 2025
